# Distinct modes of holobiont specialization among cryptic coral lineages

**DOI:** 10.1101/2024.07.05.598868

**Authors:** Carsten G.B. Grupstra, Kirstin S. Meyer-Kaiser, Matthew-James Bennett, Maikani O. Andres, David J. Juszkiewicz, James E. Fifer, Jeric P. Da-Anoy, Kelly Gomez-Campo, Isabel Martinez-Rugerio, Hannah E. Aichelman, Alexa K. Huzar, Annabel M. Hughes, Hanny E. Rivera, Sarah W. Davies

**Affiliations:** Department of Biology, Boston University, Boston, MA, USA; Biology Department, Woods Hole Oceanographic Institution, Woods Hole, MA, USA; MARE, Guia Marine Laboratory, Faculty of Sciences, University of Lisbon, Cascais 2750-374, Portugal; Palau International Coral Reef Center, Koror, Palau 96940; Coral Conservation and Research Group (CORE), Trace and Environmental DNA Laboratory (TrEnD), School of Molecular and Life Sciences, Curtin University, Bentley, Western Australia 6102, Australia; Department of Ecology, Evolution, and Marine Biology, University of California San Diego; Helmholtz-Institute for Functional Marine Biodiversity at the University of Oldenburg (HIFMB), 26129 Oldenburg, Germany; Alfred Wegener Institute, Helmholtz-Centre for Polar and Marine Research (AWI), Bremerhaven, Germany; Institute for Chemistry and Biology of the Marine Environment (ICBM), Carl von Ossietzky Universität Oldenburg, 26129 Oldenburg, Germany; Department of Integrative Biology, University of Texas at Austin, Austin, TX, USA; Northeastern University Marine Science Center, Nahant, MA, USA

## Abstract

As ocean warming threatens reefs worldwide, identifying corals with adaptations to higher temperatures is critical for conservation. Genetically distinct but morphologically similar (*i.e.,* cryptic) coral populations can be specialized to extreme habitats and thrive under stressful conditions. These corals often associate with locally beneficial microbiota (Symbiodiniaceae photobionts and bacteria), clouding interpretation of the drivers of thermal tolerance. Here, we leverage a holobiont (massive *Porites*) with high host-partner fidelity to investigate adaptive variation across classic (“typical” conditions) and extreme reefs characterized by higher temperatures and light attenuation. We uncovered three cryptic lineages that exhibit limited micro-morphological variation; one lineage dominated classic reefs (L1), one had more even distributions (L2), and a third was restricted to extreme reefs (L3). Two lineages were more closely related to populations ∼4300 km away, suggesting that these lineages are widespread. All corals harbored *Cladocopium* C15 photobionts, but strain-level compositions differed among lineages and reef types. L1 associated with distinct photobionts and bacteria in each reef type, whereas L2 had relatively stable associations. L3 hosted unique photobiont strains, signaling high host-photobiont fidelity. Analysis of light harvesting capacity and thermal tolerance revealed key adaptive variation underpinning survival in distinct habitats. L1 had the highest light absorption efficiency and lowest thermal tolerance, suggesting it is a classic reef specialist. L3 had the lowest light absorption efficiency and the highest thermal tolerance, showing that it is an extreme reef specialist. L2 had intermediate light absorption efficiency and thermal tolerance, signaling habitat generalism, potentially explaining how it survives well in both habitat types. These findings reveal diverging holobiont strategies to cope with extreme conditions. Resolving coral lineages is key to understanding variation in thermal tolerance among coral populations; uncovering thermally-tolerant holobionts can strengthen our understanding of coral evolution and symbiosis, and support global conservation and restoration efforts.

## Introduction

Corals are holobionts composed of host animals and diverse microbiota, including populations of photobionts in the family Symbiodiniaceae that provide energy from photosynthesis. Other microbiome members (bacteria, archaea, fungi, viruses and various protists) also provide a range of services including nutrient cycling and protection from pathogens (Mohamed et al., 2023). Increasing temperatures associated with global climate change can disrupt partnerships between corals and their photobionts by compromising photosynthetic activity, which can trigger photobiont loss (Baird et al., 2009; Weis, 2008) and disruption of bacterial communities (Vompe et al., 2024; Voolstra et al., 2024). The resultant nutritionally compromised “bleached” state can ultimately lead to mortality (Hughes et al., 2017). Coral bleaching events have become increasingly frequent and severe due to increased greenhouse gas emissions and local stressors (Donovan et al., 2020; Hughes et al., 2017), leading to the loss of 14% of coral reefs worldwide in under a decade (Souter et al., 2021). These events raise questions about whether coral reefs will persist in the future (Klein et al., 2024), and if so, which adaptations and symbioses will facilitate survival.

Extreme coral reef habitats characterized by naturally higher mean temperatures (Schoepf et al., 2023) can serve as a space-for-time substitution to understand coral reef futures. These reefs can have higher daily mean temperatures than nearby classic reefs, yet they maintain relatively high coral cover. Extreme reefs select for coral genotypes and microbiota that are able to persist in warmer waters, thereby acting as natural laboratories and providing potential for genetic rescue (Gonzalez et al., 2013). Some broad-level coral life history traits (*e.g.,* growth forms, growth rates) are known drivers of coral thermal tolerance (Darling et al., 2012). Additionally, given that photosynthesis disruption is at the core of the bleaching response, adaptations in light harvesting traits of coral holobionts are also likely to underpin survival in high temperatures (Enríquez et al., 2017; Gómez-Campo et al., 2022; Scheufen, Iglesias-Prieto, et al., 2017; Scheufen, Krämer, et al., 2017; Swain et al., 2018). However, less is known about how variation in these traits aligns with thermal tolerance among species with similar morphologies and life histories.

Recent sequencing efforts have uncovered many examples of cryptic coral lineages (genetically distinct but morphologically similar lineages) that are structured across environmental gradients (e.g., light, temperature, reviewed in Grupstra et al., 2024). Some cryptic lineages are more abundant on extreme reefs and appear to have higher thermal tolerance than lineages inhabiting classic reefs (e.g., Rivera et al., 2022; Rose et al., 2021; M. J. Van Oppen et al., 2018), raising questions regarding the traits that underpin variation in thermal tolerance in such closely related and morphologically similar taxa. Cryptic lineages are also often associated with genetically distinct photobiont strains, species, or even genera, that influence thermal tolerance (e.g., Durusdinium, Rose et al., 2021; Palacio-Castro et al., 2023; also see Johnston et al., 2022; Starko et al., 2023), further complicating predictions of lineage responses to climate change.

The coral genus *Porites* Link, 1807 is particularly well-suited for understanding the traits that underpin variation in thermal tolerance among cryptic lineages. Massive *Porites* corals (including morphospecies *Porites australiensis*, *P. lobata*, and *P. lutea*) are major reef-builders across the Pacific Ocean, and they inhabit classic and extreme reefs. Massive *Porites* exhibit high fidelity for *Cladocopium* C15 photobionts via vertical transmission (Bennett et al., in review; Forsman et al., 2020). Due to this tight partnership, variation in thermal tolerance among lineages is therefore likely the result of local adaptation in host and strain-level variation in photobiont associations, as well as interactions between other important holobiont members, such as bacteria (e.g., Ziegler et al., 2017). Bacterial communities of massive *Porites* are generally dominated by members of the family Endozoicomonadaceae, which can provide a variety of services to the coral holobiont, including nutritional benefits (Fifer et al., 2022; Pogoreutz & Ziegler, 2024; Vompe et al., 2024). However, high temperatures can disrupt partnerships with Endozoicomonadaceae, leading to shifts to less beneficial microbial communities (Vompe et al., 2024; but see Hadaidi et al., 2017).

Taxonomy of massive *Porites* is challenged by a lack of diagnostic characters to differentiate species. The genus contains several clades, each harboring one or more morphotypes (Forsman et al., 2009; Terraneo et al., 2021). Some morphotypes are also represented in multiple genetic clades, necessitating molecular tools to differentiate species or lineages (Forsman et al., 2009). Thus far, diverse genetically distinct, apparently cryptic, lineages of massive Porites have been identified throughout the Pacific basin (Afiq-Rosli et al., 2021; Boulay et al., 2014; Rivera et al., 2022; Starko et al., 2023; Tisthammer et al., 2020). Some of these lineages appear to be structured along environmental gradients (Boulay et al., 2014, p. 20; Rivera et al., 2022; Tisthammer et al., 2020) and several differ in terms of thermal tolerance (Boulay et al., 2014; Rivera et al., 2022; Starko et al., 2023; but see Forsman et al., 2020). However, it remains unclear whether these lineages represent widely-distributed described species, or whether they are localized taxa. It is also unknown to what extent distinct holobiont traits and symbiotic partnerships contribute to thermal tolerance.

Here, we identified colonies of three cryptic lineages of massive *Porites* that exhibit heterogeneous distributions across extreme and classic reef sites in the Rock Island Southern Lagoon of Palau. Using these lineages, we aimed to answer five critical questions: 1) Do these lineages exhibit micromorphological differences that are consistent within current *Porites* taxonomy? 2) To what extent are they related to *Porites* lineages recently discovered elsewhere in the Pacific Ocean? 3) Do these lineages exhibit distinct symbioses? 4) Are these lineages functionally distinct? 5) How do holobiont interactions shape thermal tolerance? Understanding how holobiont symbioses and physiology interact to shape cryptic lineage distributions and thermotolerance is key to understanding how coral reefs will respond to future climate change.

## Methods

### Software

All data were analyzed in R v4.1.2 (R Core Team, 2019). Linear models (LM) were conducted using the lm function in the package stats 4.1.2, linear mixed effects models (LMM) with lmer function in lme4 v1.1-31 (Bates et al., 2014); F-tests in car v3.0-12 were used for the selection of significant variables (Fox et al., 2019). Post-hoc pairwise comparisons were conducted using emmeans v1.8.4-1 (Lenth et al., 2023). We assessed model assumptions using performance v0.10.2 (Lüdecke et al., 2021). Permutational multivariate ANOVAs (PERMANOVAs) were conducted using the adonis function in vegan 2.6-4 (Oksanen et al., 2019).

### Site selection and sample collection

Three extreme and three paired classic sites with similar depth (2 - 4 m) and proximity to land were selected in Chelbacheb (The Rock Islands of Palau, Figure 1A). Extreme sites are semi-enclosed lagoons with higher mean water temperatures, increased light attenuation, and distinct assemblages of massive *Porites* lineages compared to classic sites (Rivera et al., 2022; van Woesik et al., 2012). Water temperatures were measured at each site between November 2021 and May 2022, and light levels were measured for 16 days in April 2022 (Onset, Wareham, USA). Colonies resembling the gross morphology of *Porites lobata* Dana, 1846 were tagged at all sites in November 2021 and tissue samples were taken from the center of each colony and immediately stored in ethanol at -20℃ (2×2 cm samples; *n* = 90 total, 15/site, Table S1). An additional 22 colonies were sampled in April 2022 (*n* = 2-7 per site, Table S1). We tested for differences in temperature (daily mean, maximum, minimum, and range) and light intensity levels (mean, maximum, range) between reef types (classic, extreme) using an LMM with individual sites included as a random effect.

**Figure 1.**
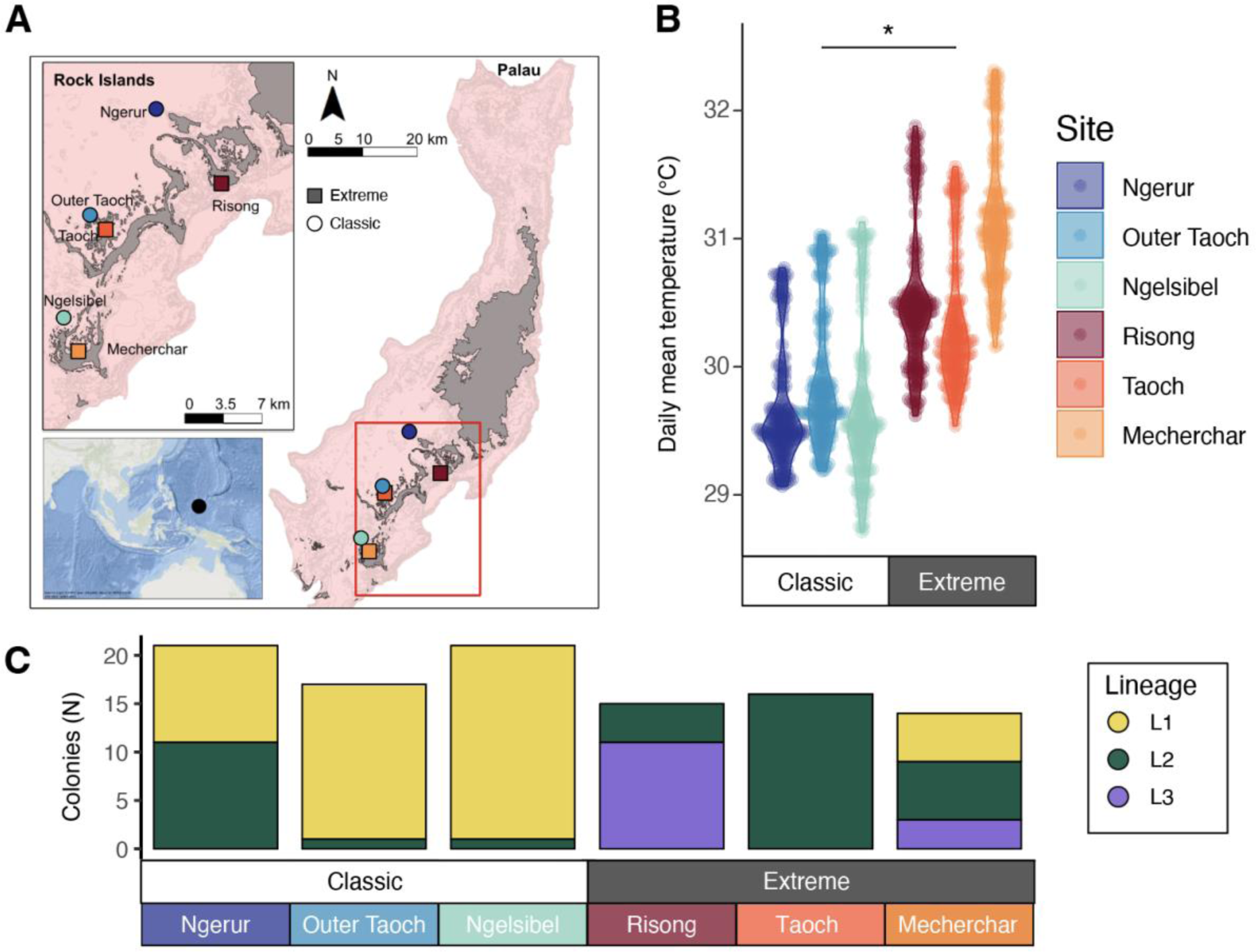
Collection sites, temperature conditions, and distributions of three cryptic massive *Porites* lineages at six study sites in Palau, Micronesia. A) Three paired sites were selected (*n* = 6 sites total), with each pair consisting of one exposed site (“classic”, circles) and one enclosed lagoon site (“extreme”, squares). B) Daily mean temperatures between November 2021 and April 2022 were on average ∼1 ℃ higher at extreme sites (30.7℃ ± 0.64) relative to classic sites (29.7℃ ± 0.54, *F* = 13.25, *p* = 0.02). C) Relative abundances of L1 (yellow) colonies were higher on classic reefs, whereas L2 (green) colonies had relatively even distributions among reef types; L3 (purple) colonies were restricted to extreme reefs. Lineages were identified *via* 2b-RAD sequencing across two sequencing runs (*n* =104 total, Table S1). See Figure S3, S4 for additional detail on lineage classifications.

### Coral genetics and micro-morphological observations

DNA was extracted from all samples and libraries were generated for population genetics approaches using 2b-RAD sequencing (Rippe et al., 2021; Wang et al., 2012) (see Supplementary Methods). Note that 2b-RAD sequencing was conducted in two runs, but all population genetics metrics (F_st_, admixture, principal component analysis) were generated based on 75 samples collected in November 2021 (out of 90) that successfully sequenced in the first run. This dataset was then reanalyzed in combination with another recent 2b-RAD dataset that identified cryptic lineages in Kiritimati to determine whether these represent different populations of the same lineages (Starko et al., 2023). Samples collected in Palau in April 2022 (*n* = 22), as well as failed libraries from November 2021 (*n* = 15), were sequenced in a later run using a reduced representation design and used for lineage assignment only (overall successful 2b-RAD sequencing *n* = 104/112; Table S1). The combined data from the two sequencing runs were used to test the hypothesis that lineages differ in terms of their distributions across classic and extreme sites using a Chi-squared test. Then, micro-morphological analysis of skeletal fragments from Palau (*n =* 19) was conducted to test whether the lineages were morphologically similar to described species (see Supplementary Methods).

### Characterization of microbial communities

Characterization of photobiont and bacterial communities associated with each lineage was conducted on samples collected in November 2021 using ITS-2 and 16S sequencing, respectively (see Supplementary Methods; Table S1). Raw ITS-2 reads were processed by Symportal (Hume et al., 2019) to produce defining intragenomic sequence variant (DIV) profiles for each coral colony (final *n* = 73; Table S1). All colonies were dominated by C15 DIVs. cumulative link models (clm) were used to test for differences in the dominant C15 DIV associations by including an interaction between lineages and reef type. Site was also included as an additional factor (clm does not allow random or nested factors) in the model to account for local variation.

Quality filtering, denoising, merging, and taxonomy assignments of 16S rRNA gene reads were conducted with DADA2 against the *Silva* v. 138.1 database (Quast et al., 2012), and contaminant reads were removed in Phyloseq (McMurdie & Holmes, 2013) (final *n* = 48; see Table S1, Supplementary Methods). We tested for differences in bacterial diversity (Shannon and Simpson indices, calculated based on the non-rarefied dataset using the estimate_richness function) using an LM with an interaction between lineage and reef type. Including site as a random effect resulted in model convergence failure, so it was removed. Bray-Curtis distances were calculated based on data from which rare (<10 reads) ASVs were removed (2361 remaining taxa). We tested for differences in bacterial community composition using a PERMANOVA with an interaction between lineage and reef type and collection site was nested within reef type.

### Analyses of holobiont optical traits

To characterize differences in holobiont structural and optical traits, we quantified polyp densities and light-harvesting characteristics in a subset of coral colonies (Table S1; Supplementary Methods). A total of 20 tagged colonies of known lineage and reef type (Table S1) were transported to Boston University in May 2023, fragmented, and then acclimated for 63 days in aquariums. Reflectance (R) between 400 and 750 nm was measured (sensu Enríquez et al., 2005; Vásquez-Elizondo et al., 2017) (see Supplementary Methods). Chlorophyll *a* densities were quantified and the specific absorption coefficient of Chlrophyll *a* (Chl*a*; a*_chla_), a measure of the light absorption efficiency of the holobiont, was estimated following Enríquez et al., (2005) (see Supplementary Methods). We tested for differences in chlorophyll concentration and a*_chla_ values among lineages using an LM with reef type and lineage as fixed effects (no interaction because we only had L2 colonies from both reef types). Pairwise comparisons among lineages were conducted with a Bonferroni *p*-value correction.

### Thermal challenge experiment

A 25-day common garden heat challenge experiment was conducted to test for differences in thermal tolerance between the three *Porites* lineages (*n* = 24; Table S1; see Supplementary Methods). L1 colonies were collected from classic sites and L2 and L3 colonies were collected from extreme sites. We initially aimed to include L1 and L2 colonies from both extreme and classic sites, but were unsuccessful. Two cores were extracted from each colony; one core was assigned to the control treatment, and the other to the heat treatment. Mean temperatures in control tanks (*n* = 3 tanks per treatment) were maintained at 29.5°C ± 0.1°C. Temperatures in the heat treatment tanks were ramped by ∼3°C over seven days, followed by a 12-day hold. On day 19, temperatures were increased by an additional ∼1°C to simulate an extreme thermal stress event until day 25. All cores were inspected daily for mortality. Maximum PSII photochemical efficiency (Fv/Fm) was measured daily or semi-daily following >90 minutes of dark incubation. All fragments were also photographed with a color standard at six timepoints to measure changes in coloration (paling), quantified as the intensity in the gray channel using ImageJ (sensu McLachlan & Grottoli, 2021). To control for differences in gray intensity at the start of the experiment, we also calculated and analyzed relative changes in gray intensity (Δ grey intensity; values in heat - values in control for each colony at each timepoint).

Lineage-specific survival rates were estimated with the Kaplan-Meier method and a G-test using the packages survival v3.3-1 (Therneau, 1999) and survminer v0.4.9 (Kassambra, 2018). To test how lineage and thermal challenge affected Fv/Fm and bleaching of coral colonies over time, we used an LMM with a three-way interaction between treatment, lineage, and time. Colony number was included as a random effect to account for repeated sampling. Photobiont DIV was not included in the models because it co-varied with lineage. For relative coloration, the same analysis was done but treatment was not included in the analysis (because the values for heat-treated corals were relative to controls). Post-hoc pairwise comparisons between lineages for each day across treatments were conducted with a false discovery rate (Fv/Fm and grey intensity) or Bonferroni (Δ grey intensity) correction.

## Results

### Extreme sites are characterized by higher mean temperatures

Daily mean, maximum, and minimum temperatures were 0.8-1.1℃ higher at extreme sites (Figure 1B, S1, Supplementary Datafile 1) than at paired classic sites (LMM results: *F* = 13.25, *p* = 0.02; *F* = 25.88, *p* = 0.01; *F* = 9.08, *p* = 0.04). Daily temperature range did not differ between extreme and classic sites (Figure S1; Supplementary Datafile 2). Light intensities (mean, maximum, range) also did not differ between reef types over the 16-day measurement period (Figure S2).

### Uneven distributions of lineages between reef types

SNP data revealed three distinct genetic lineages (henceforth L1, L2, L3) in Palau (Figure 1C, S3, S4). Including only samples collected in 2021 (Table S1), weighted F_ST_ values using all loci (487,248 loci) were 0.286 between L1 (yellow) and L2 (green), 0.468 between L1 and L3 (purple), and 0.456 between L1 and L3. When only neutral loci were included (outliers removed), F_ST_ values dropped to 0.118 between L1 and L2 (based on 844,221 loci, after 5,882 outlier loci were removed), 0.289 between L1 and L3 (purple; 765,683 loci, 5,521 loci removed), and 0.281 between L2 and L3 (793,557 loci, 5,797 loci removed). Relative abundances of the three lineages (including samples sequenced in a subsequent run, *n* = 29; Table S1) differed between classic and extreme sites (*χ*^2^ = 50.32, *p* < 0.001). L1 colonies were more abundant at classic sites, while L2 exhibited more even distributions with higher abundances at extreme sites. L3 was restricted to extreme sites (Risong, Mecherchar; Figure 1C).

### Genetic connectivity is lower among co-occurring than distant lineages

A re-analysis of SNP data from Palauan samples from 2021 and data from Starko et al., (2023) showed that L1 and L2 are less genetically differentiated from lineages in Kiritimati than they are from co-occurring lineages in Palau (Figure S5; based on 13443 loci). Specifically, L1 was more genetically similar to a lineage in Kiritimati (Pkir-2, *F_ST_* = 0.16) than to L2 or L3 (*F_ST_* = 0.24-0.42). Additionally, L2 was more genetically similar to another distinct lineage in Kiritimati (Pkir-1, *F_ST_* = 0.22) than to L1 or L3 (*F_ST_* = 0.24-0.41). L3 was highly differentiated from all other lineages, regardless of location (*F_ST_* = 0.41-0.52). A third Kiritimatian lineage (Pkir-3) was also highly diverged from all other lineages (*F_ST_* = 0.38-0.52), potentially indicating endemic lineages at both locations. Regardless, admixture analysis suggested that lineages in Palau and Kiritimati likely all represent reproductively isolated populations (Figure S6).

### Limited micro-morphological traits differentiate some lineages

Micro-morphological characterisation from Z-stacked photos and SEM images from 19 coral fragments revealed morphological characters differentiating some lineages (Figure 2, see Supplementary Results). However, due to morphological overlap among the three lineages, they are challenging to distinguish based on morphology alone. A key distinctive feature of L1 (Figure 2A-D) is the occasional fusion of the ventral directive to the lateral septa of the triplet, resulting in a large palus almost equal in size to the pali of the lateral pairs (Figure 2D). The morphological characteristics resemble those of *Porites australiensis* Vaughan, 1918, with connotations of *Porites lutea* Milne Edwards & Haime, 1851. However, the absence of frequent triplet fusion or trident formation suggests a closer affinity with the former.

**Figure 2.**
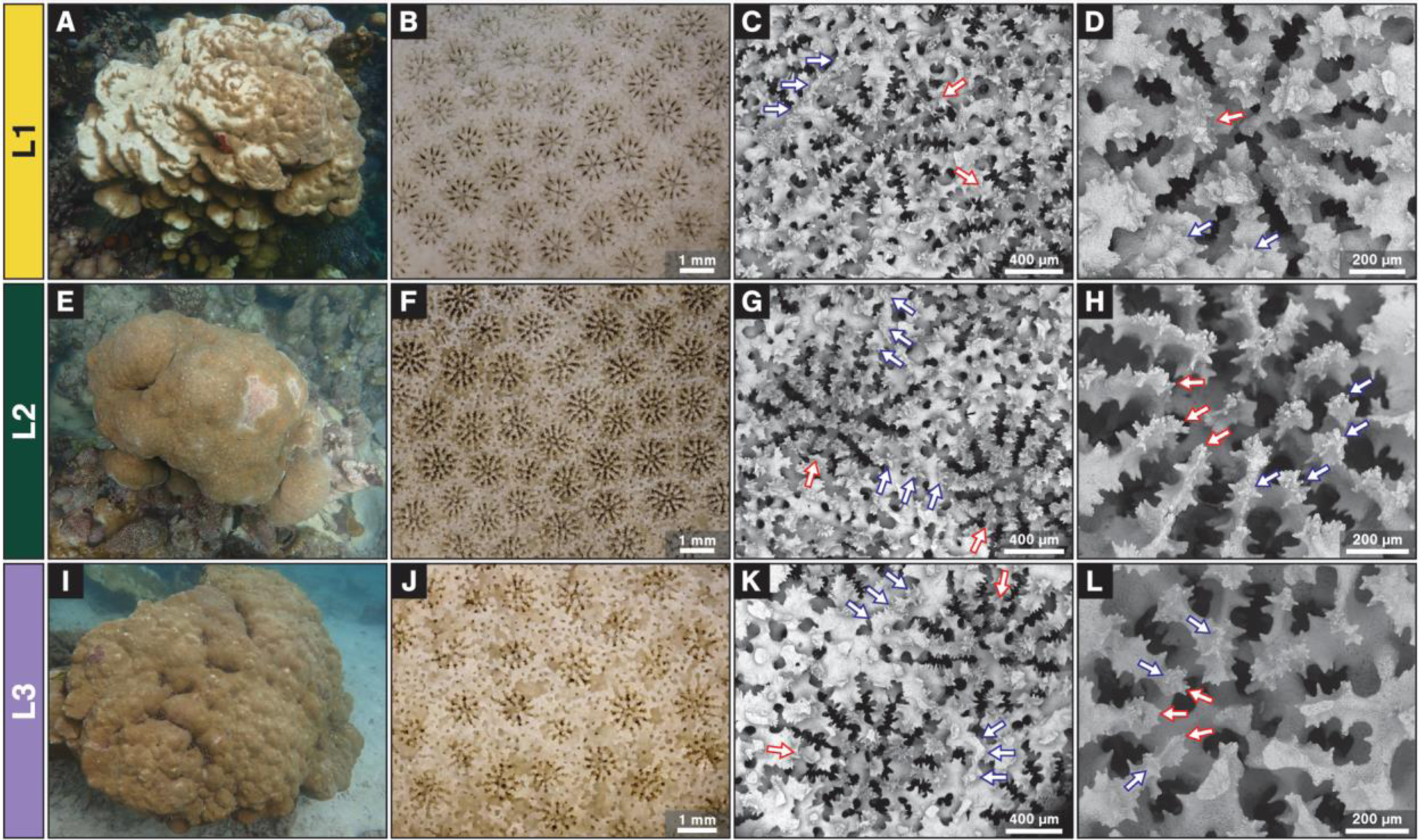
*In situ* colonies and corallite morphology of *Porites* specimens collected from Palau, Micronesia, consisting of various representatives of L1 (A–D), L2 (E–H) and L3 (I–L). A) Large colony with a series of thick ledges; B) Corallites displaying thick straight walls and thick septa; C) Corallites with a free ventral triplet (white-red-outline arrows) and ridge-like walls with rough mural denticles (white-blue outline arrows); D) Corallite with fusion of the ventral triplet, forming a large single palus (white-red-outline arrow), with inner denticles smaller in size than the corresponding palus of the lateral pair septa (white-blue-outline arrows); E) Medium sized hemispherical colony with a smooth appearance; F) Corallites displaying moderately excavated calices with medium thick walls; G) Corallites with a free ventral triplet (white-red-outline arrows) and walls composed of three rows of denticles (white-blue outline arrows); H) Corallite with free ventral triplet (white-red-outline arrows) and inner denticles slightly more prominent or equal to the size of its corresponding septal palus (white-blue-outline arrows); I) Large hemispherical colony with a hillocky appearance; J) Colony displaying moderately excavated calices with thick walls; K) Corallites with a free ventral triplet (white-red-outline arrows) and a ridge-like wall with rough granulated mural denticles (white-blue outline arrows); L) Corallite with free ventral triplet (white-red-outline arrows) and poorly developed pali smaller or equal to its corresponding septal denticle (white-blue-outline arrows). (A, E, I) *In situ* colony images from Risong, depth 2-4 meters; (B, F, J) Z-Stacked light microscope images; (C, D, G, H, K, L): SEM images of corallum surface.

L2 (Figure 2E-H) and L3 (Figure 2I-L) can be distinguished from L1 because they lack fusion of the ventral directive triplet observed in L1. Yet, they are challenging to distinguish from each other based on microskeletal features. The morphological characteristics of L2 and L3 both resemble those of *Porites lobata* Dana, 1846. Yet, L3 shares some morphological similarities with L1 and *P*. *australiensis*, such as reduced length of the dorsal and ventral septum and well-formed pali usually larger or equal to the septal denticles.

### Photobiont associations differ among lineages and reef types

All sampled colonies were dominated by *Cladocopium* C15 with 10 unique C15 DIVs identified (Figure 3; Supplementary Datafile 3). Photobiont associations of L1 colonies differed between classic and extreme reefs: colonies at classic sites (*n =* 31) hosted a variety of DIVs, while extreme L1 colonies all harbored C15.C93a (*n* = 4). By comparison, patterns of photobiont associations in L2 were more similar between classic and extreme sites: colonies at both reef types hosted diverse DIVs, three of which were shared between classic (out of 4 DIVs) and extreme sites (out of 5 DIVs). Most (*n* = 8/10) L3 colonies hosted unique C15 DIVs (C15.C15vp and C15.C15vp.C15vt) that were not found in L1 or L2.

**Figure 3.**
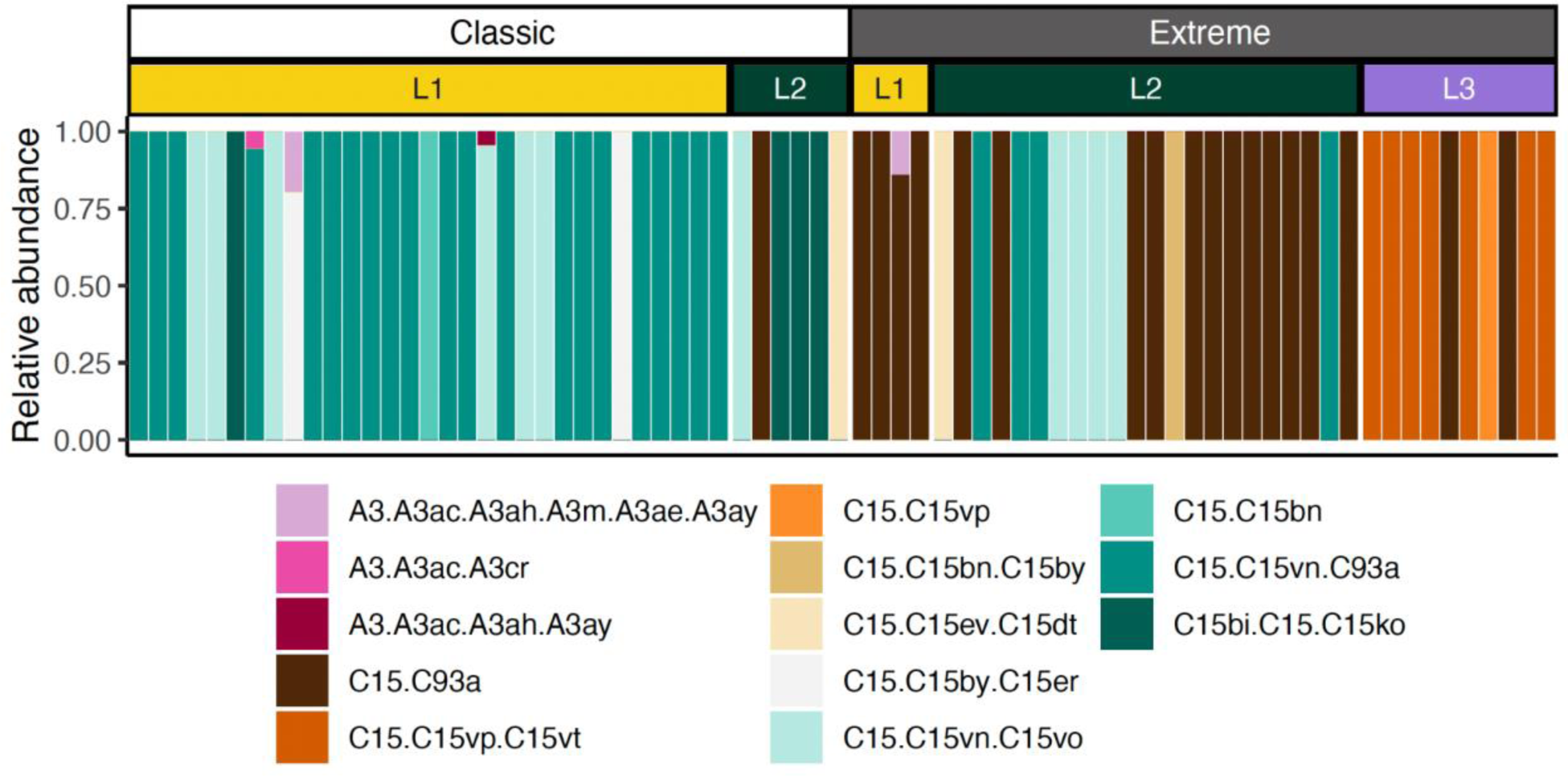
Relative abundances of photobiont (Symbiodiniaceae) defining intragenomic variants (DIVs) across all three *Porites* lineages in classic and extreme reef types. All colonies were dominated by *Cladocopium* C15 DIVs. Note that one DIV is present in 50% of extreme reef colonies (C15.C93a, *n* = 18/36), while it was only found in 1 (out of 37) classic reef colony. Additionally, most (*n* = 8/10) L3 colonies hosted unique C15 DIVs (C15.C15vp and C15.C15vp.C15vt) not found in L1 or L2. Sample sizes (classic, extreme sites): L1 (31, 4); L2 (6, 22); L3 (0, 10).

Overall, C15.C15vn.C93a was the most abundant DIV, which was identified in 21/31 L1 and 4/28 L2 colonies and dominated L1 colonies at classic sites. One DIV present in 50% of extreme site colonies (C15.C93a, *n* = 18/36) was only found in one (out of 37) classic reef colony. Cumulative link model results showed that reef type (*Df* = 1, *χ^2^* = 60.3, p<0.001), site (*Df* = 5, *χ^2^* = 57.7, *p <* 0.001), and lineage (*Df* = 2, *χ^2^* = 56.1, p<0.001) had similarly sized effects on photobiont associations. There was also a significant interaction between reef type and lineage (*Df* = 2, *χ^2^* = 6.8, *p* < 0.03), indicating that photobiont populations differ among reef types in lineage-specific ways. Of note, four L1 colonies (*n =* 3 classic, 1 extreme) had low abundances of *Symbiodinium* A3 (< 25% relative read abundance).

### Bacterial communities differ among lineages and reef types

Bacterial (alpha) diversity was higher in colonies sampled at extreme sites than at classic sites but did not differ between lineages (Figure 4A, B; Simpson LM results: Lineage *Df* = 1, *F* = 0.31, *p* = 0.73; Reef type: *Df* = 1, *F* = 7.11, *p* = 0.01, Lineage*Reef type: *Df* = 1, *F* = 3.13, *p* = 0.084; Shannon LM results: Lineage *Df* = 2, *F* = 0.21, *p* = 0.81; Reef type: *Df* = 1, *F* = 8.48, *p* = 0.006, Lineage*Reef type: *Df* = 1, *F* = 0.27, *p* = 0.61; Supplementary Datafile 4). Specifically, mean (±SE) Simpson values were 1.17 (L2) - 2.1 (L1) times higher (classic: L1 = 0.45 ± 0.06 L2 = 0.65 ± 0.04; extreme: L1 = 0.93 ± 0.04, L2 = 0.76 ± 0.06, L3 = 0.85 ± 0.08) and Shannon index values were 2.00 (L2) - 2.3 (L1) times higher at extreme sites than classic sites (classic: L1 = 1.61 ± 0.29, L2 = 1.52 ± 0.16; extreme: L1 = 3.70 ± 0.53, L2 = 2.99 ± 0.56, L3 = 2.86 ± 0.65).

**Figure 4.**
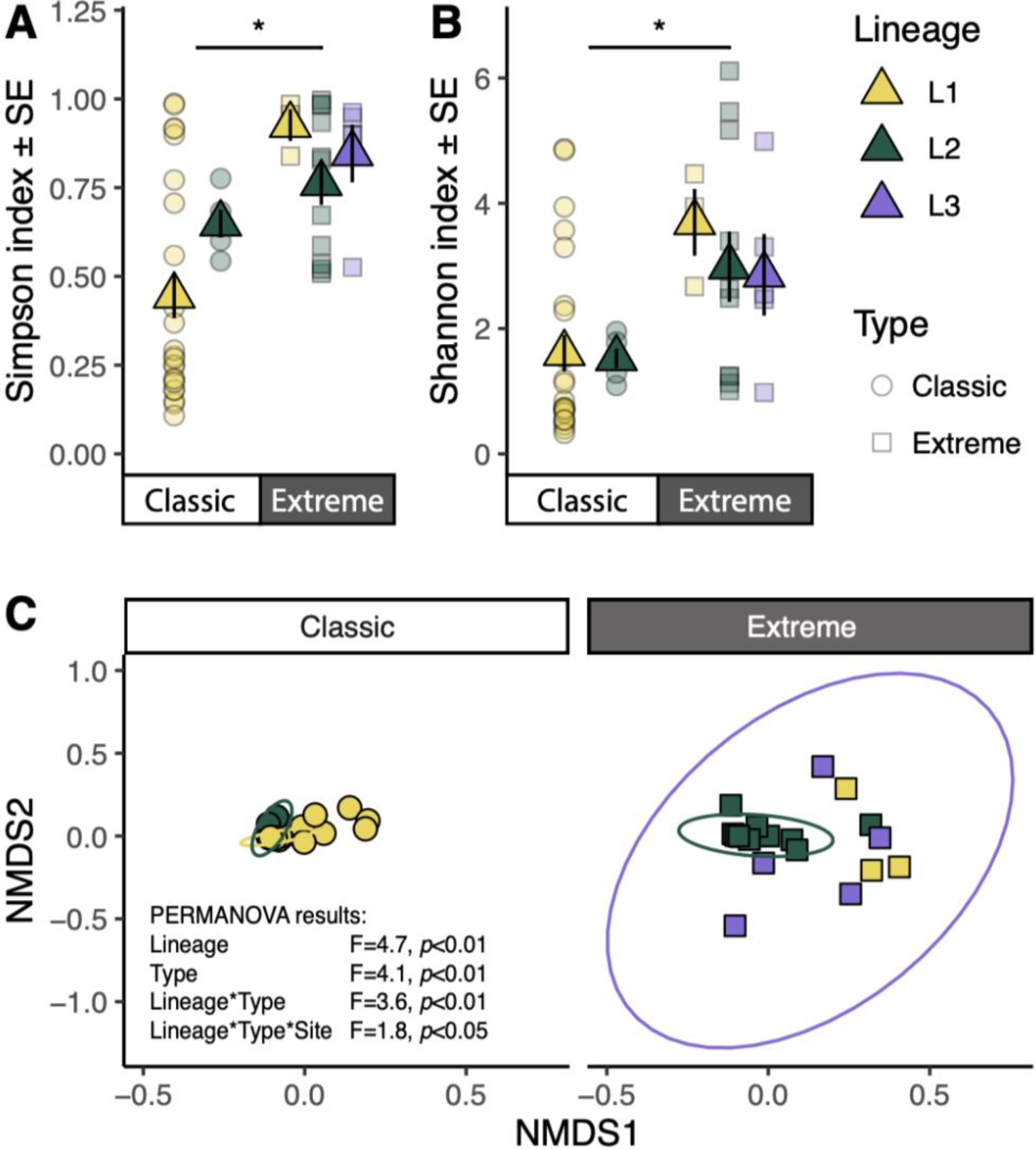
Bacterial community compositions differ between reef types and among lineages of massive *Porites.* Mean Simpson (A) and Shannon (B) metrics of bacterial diversity were significantly higher in corals sampled at extreme sites than classic sites. C) Non-Metric multidimensional scaling (NMDS) plots reveal differences in bacterial community compositions among lineages, reef types, and sites. NMDS stress = 0.15. Ellipses denote 95% confidence interval. Sample sizes (classic, extreme sites): L1 (24, 3); L2 (5, 11); L3 (0, 5).

Coral lineage was the most important driver of bacterial community compositions (Figure 4C; *F* = 4.7, *R^2^* = 0.15, *p* = 0.001), followed by reef type (*F* = 4.1, *R^2^* = 0.07, *p* = 0.002). There were also significant interactions between lineage and reef type (*F* = 3.55, *R^2^* = 0.06, *p* = 0.001), and between lineage, reef type, and site (*F* = 1.8, *R^2^* = 0.11, *p* = 0.016). Dispersion was not significantly different among sampling groups (*Df* = 4, *F* = 1.92, *p* = 0.12).

L1 and L2 colonies at classic sites were dominated by Endozoicomonadaceae (Figure 5A). At extreme sites, only L2 colonies harbored high relative Endozoicomonadaceae abundances. No L3 colonies hosted appreciable proportions of this genus. Associations with particular Endozoicomonadaceae at the ASV level also differed among lineages (Figure 5B): L1 colonies at classic sites were dominated by ASV1, with many colonies also hosting ASV 6 ( *n* = 15/24) and 12 (*n* = 14/24). L2 colonies at classic sites also all hosted ASVs 1 and 6, but no colonies at extreme sites hosted ASV 6. Most L2 colonies hosted ASV 10 instead of 12 ( *n* = 14/16), regardless of reef type, and half of L2 colonies at extreme sites also hosted ASV 30 (*n* = 5/10). L1 and L3 colonies at extreme sites all hosted low abundances of ASV 1 (*n* = 8/8).

**Figure 5.**
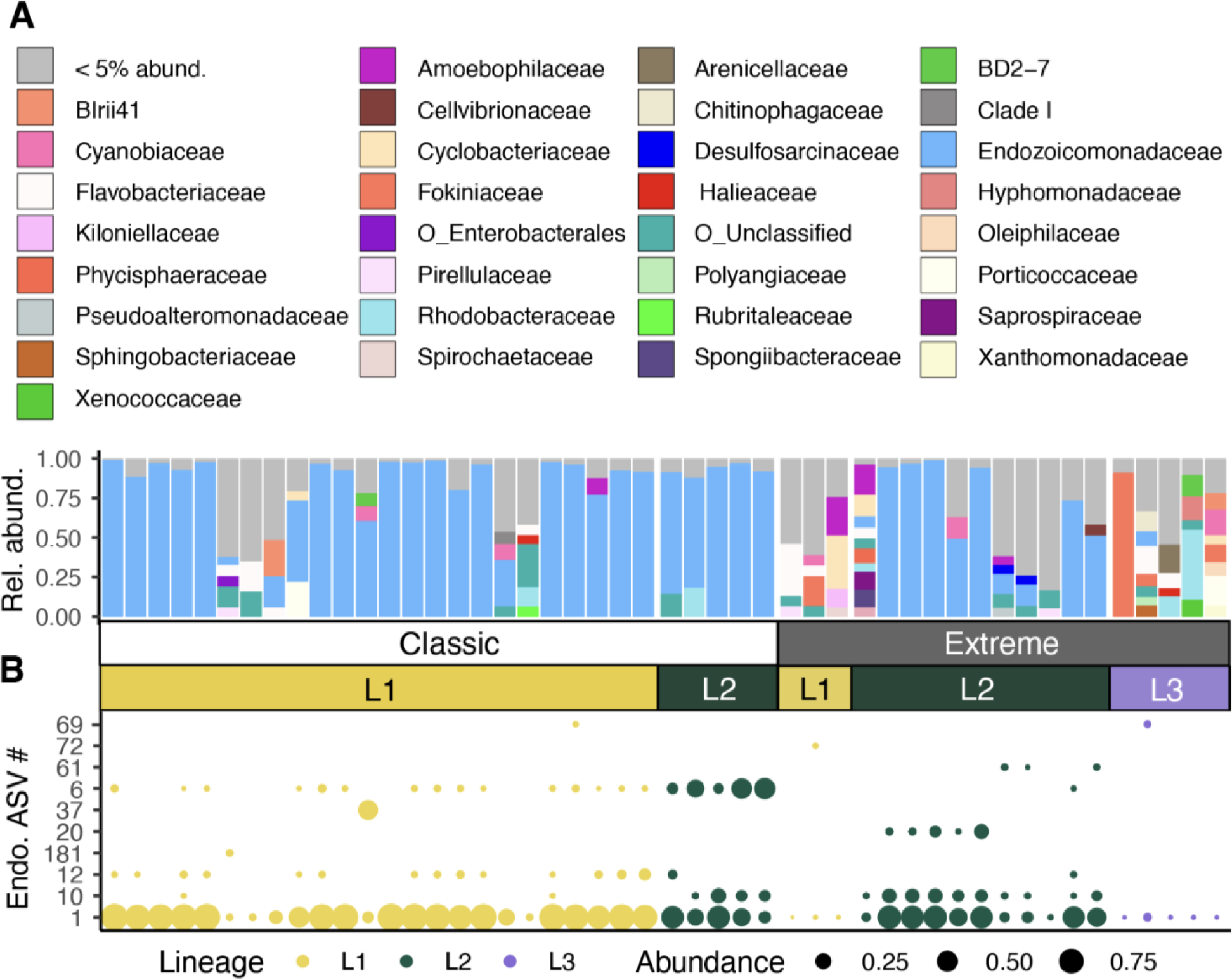
Bacterial community composition patterns differ between lineages of massive *Porites*. A) Relative abundances of bacterial families across samples demonstrating that taxonomic compositions differed between lineages and reef types (Classic, Extreme). Rel. abund.=Relative abundance. B) Relative abundances of top 10 most abundant Endozoicomonadaceae ASVs. Endo. ASV#=Endozoicomonadaceae amplicon sequence variant number. Sample sizes (classic, extreme sites): L1 (24, 3); L2 (5, 11); L3 (0, 5).

### Optical traits differ among lineages

Lineages differed in terms of polyp densities (Figure 6A, S7, S8; see Supplementary Results, Supplementary Datafile 5) and optical traits (Figure 6B-D; Supplementary Datafile 6). Specifically, Chl*a* densities (mean µg cm^-2^ ± SE) varied among lineages (Figure 6B; *Df* = 2, *F* = 10.65, *p* = 0.001) but not reef type (*Df* = 1, *F* = 1.30, *p* = 0.27). Pairwise comparisons (L1-L2 est = -0.292, *Df* = 16, *p* = 0.0233; L1 - L3 est = -0.594, *Df* = 16, *p* = 0.0009; L2 - L3 est = -0.301, *Df* = 16 , *p* = 0.0095) showed that L3 had the highest Chl*a* concentrations (0.76 ± 0.06), followed by L2 (0.51 ± 0.04). L1 colonies had the lowest Chl*a* concentrations (0.28 ± 0.06).

**Figure 6.**
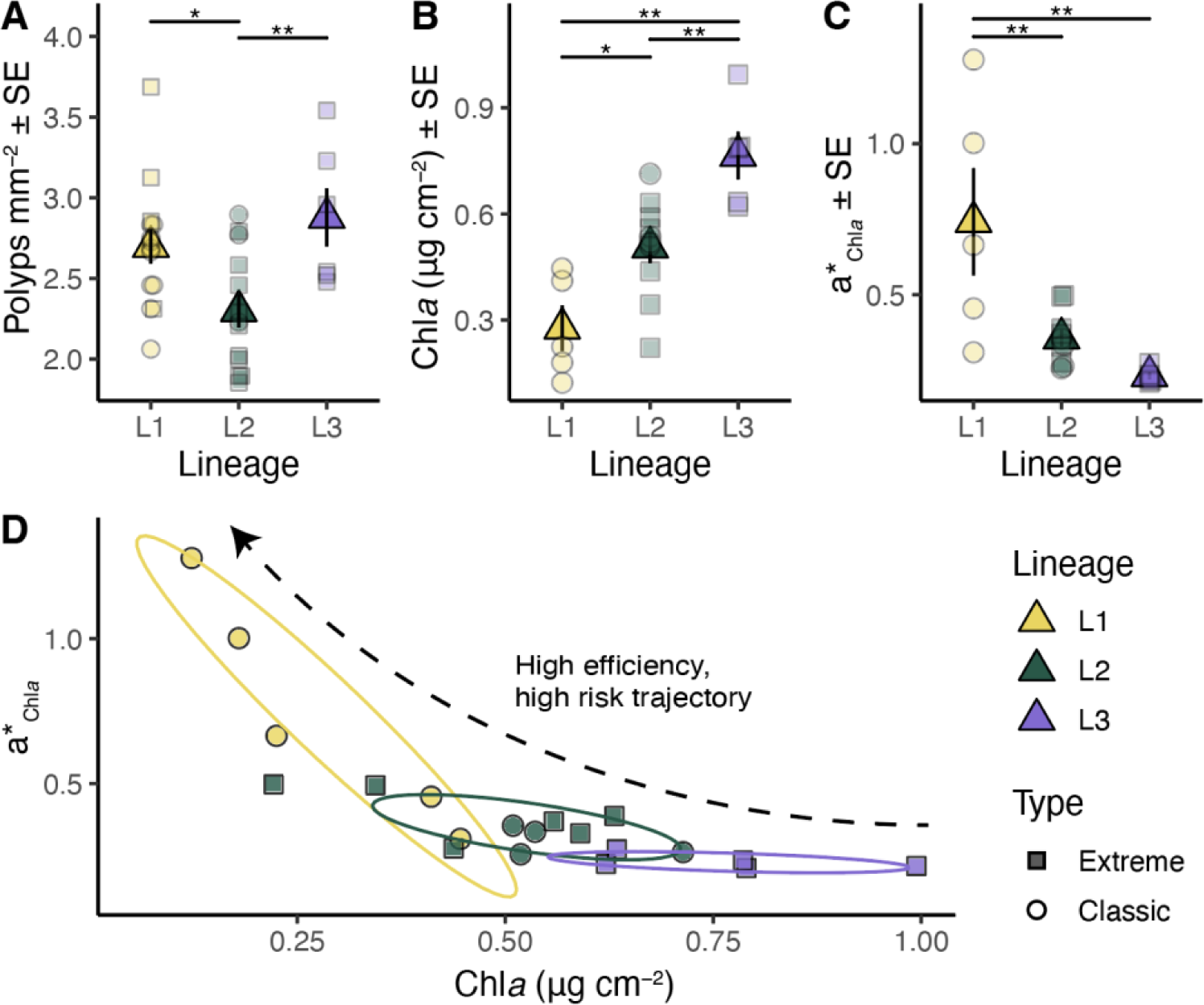
Structural and optical traits of cryptic lineages of massive *Porites* in Palau indicate functional variation and specialization to classic and extreme reef environments. A) Mean polyp densities (polyps mm^-2^) differ among lineages. B) Mean Chlorophyll *a* (Chl*a*) concentrations (µg cm^-2^) vary among lineages. C) Mean Chl*a*-specific absorption coefficient (a*_Chl*a*_, m^2^ mg chl*a*^-1^) values are higher in L1 than the other lineages. D) Higher Chl*a* absorption (a*_Chl*a*_) in L1 is correlated with lower Chl*a* concentrations, which is a “high efficiency, high risk” phenotype (sensu Scheufen, Iglesias-Prieto, et al., 2017). Ellipses in D denote 68% confidence intervals. * *p* < 0.05, ** *p* < 0.01.

Light absorption efficiency (a*_Chla_, m^2^ mg chla^-1^; Figure 6C) also varied among lineages (*Df* = 2, *F* = 5.89, *p* = 0.012) but not reef type (*Df* = 1, *F* = 0.45, *p* = 0.508). Pairwise comparisons (L1 - L2 est = 0.439, *Df* = 16, *p* = 0.0175; L1 - L3 est = 0.601, *Df* = 16, *p* = 0.0159; L2 - L3 est = 0.161, *Df* = 16, *p* = 0.6434) showed that L1 had the highest a*Chla (0.74 ± 0.18), followed by L2 (0.36 ± 0.03). L3 had the lowest a*_Chla_ (0.23 ± 0.01).

Plotting of light absorption efficiency and chlorophyll *a c*oncentrations in the same figure (Figure 6D) visually demonstrated that the three lineages differ from one another in terms of optical traits; L3 colonies corresponded to low light efficiency, low risk, phenotypes; L1 corresponded to a high light efficiency, high risk, phenotypes; and L2 was intermediate (sensu Scheufen, Iglesias-Prieto, et al., 2017).

### Variation in thermal tolerance among lineages

Heat challenge affected coral survival, photosynthetic efficiency (Fv/Fm), and coloration; however, the strength of responses to heat differed among lineages (Table 1, Figure 7). Thermal challenge (Figure 7A) caused mortality in 46% (*n* = 11/24) of coral fragments (Figure 7B; Supplementary Datafile 7). Mortality was first observed in L1 at day 12, and in L2 at day 16; mortality of L1 and L2 progressed until the end of the heat challenge. Kaplan-Meier analysis demonstrated that survival rates significantly differed between the lineages (Figure 7B; *χ*^2^ = 6, *Df* = 2, *p* = 0.049), and was lowest in L1 corals: full mortality was observed in 70% of L1 (*n* = 7/10), whereas only 44% of L2 died (*n* = 4/9). No mortality was observed in L3 (*n* = 0/5).

**Table 1.** Outputs of global linear mixed-effects models for maximum photochemical efficiency (Fv/Fm), coloration (grey intensity), and relative coloration (Δ grey intensity: Heat-Control). For all three models, coral colony (genotype) was included as a random effect.

| Dependent variable | Independent variable | *F* | *Df* | *p* |
| --- | --- | --- | --- | --- |
| *Fv/Fm* | Treatment | 780.4 | 1 | **<0.001** |
|  | Lineage | 5.8 | 2 | **0.011** |
|  | Day | 13.6 | 16 | **<0.001** |
|  | Treatment:Lineage | 15.6 | 2 | **<0.001** |
|  | Treatment:Day | 17.0 | 16 | **<0.001** |
|  | Lineage:Day | 1.3 | 32 | 0.113 |
|  | Treatment:Lineage:Day | 2.0 | 32 | **<0.001** |
| *Grey intensity* | Treatment | 241.9 | 1 | **<0.001** |
|  | Lineage | 15.5 | 2 | **<0.001** |
|  | Day | 10.6 | 5 | **<0.001** |
|  | Treatment:Lineage | 7.4 | 2 | **<0.001** |
|  | Treatment:Day | 29.3 | 5 | **<0.001** |
|  | Lineage:Day | 0.9 | 10 | 0.496 |
|  | Treatment:Lineage:Day | 0.7 | 10 | 0.743 |
| $\Delta$ Grey intensity: *H-C* | Lineage | 3.6 | 2 | **0.047** |
|  | Day | 40.5 | 5 | **<0.001** |
|  | Lineage:Day | 0.9 | 10 | 0.495 |

**Figure 7.**
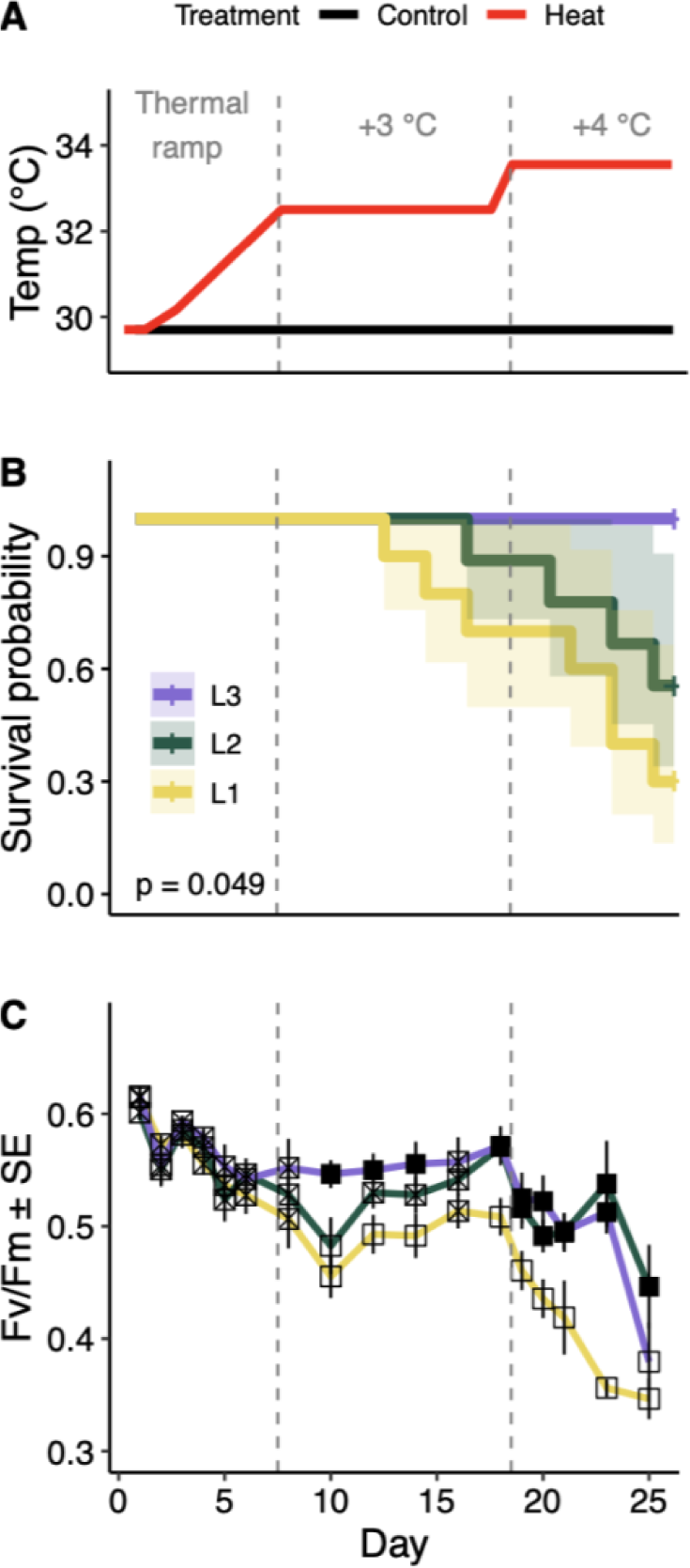
Responses to thermal challenge differed among lineages of massive *Porites.* A) Fragments of three lineages of massive *Porites* were distributed among control (∼29.5℃, *n =* 3) and heat tanks (*n =* 3) in which temperatures were raised by 3-4℃ (see Figure S9 for measured temperatures). B) Survival probability differed among lineages (± 90% CI) exposed to thermal challenge (*X^2^*= 6, *p* = 0.049). No mortality was observed in the control tanks. C) Changes in photosynthetic efficiency (Fv/Fm) in colonies exposed to thermal challenge differed among lineages. Closed points differ significantly from open points (p < 0.05; see Table S3 for pairwise test results). Hatched points did not differ significantly from other points. Sample sizes: L1 (*n* = 10; 7 died); L2 (*n* = 9; 4 died); L3 (*n* = 5; 0 died).

On day 1 of the heat challenge experiment, mean Fv/Fm of the three lineages were similar, ranging between 0.605 ± 0.008 (L3) - 0.615 ± 0.007 (L2). Heat challenge significantly reduced Fv/Fm in all lineages (Figure 7C, Table 1; Supplementary Datafile 8); however, loss of Fv/Fm across lineages was significantly different (interaction between treatment and lineage) and the rate of loss varied significantly (interaction between treatment, lineage and day). For example, Fv/Fm values were reduced after just two to four days in L1 and L2, whereas L3 was not affected until day six (Table S2). Following this initial drop in Fv/Fm after 4-6 days, lineages exhibited different responses to the continued heat stress: Fv/Fm values continued dropping for L1 and L2 until day ten of the experiment, whereas L3 remained stable and maintained significantly higher Fv/Fm values than L1 (Figure 7C; Table S3) and L2 (significant on day 10, *F* = -0.067, *p* = 0.03). Fv/Fm in L1 and L2 both showed some level of recovery between days 10 and 18, but this recovery was more pronounced for L2 (L1-L2 comparison at day 18; Figure 7C; Table S3). The compounded heat stress (day 19-25) caused further reductions in Fv/Fm values for all three lineages (Figure 7C), but L1 was more negatively affected than either of the other lineages (Table S2, S3). While not included in the model to avoid confounding effects, photobiont DIV also appeared to influence colony thermal tolerance, adding an additional layer of complexity to lineage thermal responses (Figure S10).

Analysis of changes in grey intensity from standardized photographs showed that the paling rates also differed among lineages (Figure S11; Table 1; Supplementary Datafile 9). For example, L1 colonies appeared to pale more rapidly (Figure S11, Table S4) and were significantly more paled compared to L3 colonies at day 20 (higher Δ grey intensity; Table S5).

## Discussion

Scleractinia are morphologically and functionally diverse, but genetic and functional variation are often masked by morphological plasticity or similarity among sister taxa (Bongaerts et al., 2021; Boulay et al., 2014; Burgess et al., 2021; Grupstra et al., 2024). Understanding differences in thermal tolerance among morphologically indistinguishable–cryptic–coral lineages is key to predicting coral reef fates under warming futures. Yet, this understanding is further hindered by heterogeneous associations with microbial partners that interact with host diversity to affect thermal tolerance (Palacio-Castro et al., 2023; Rose et al., 2021). Here, we provide strong evidence for three cryptic lineages (L1-L3) of a stony coral that exhibit high photobiont fidelity within the massive *Porites-Cladocopium* C15 system, in which cryptic lineages exhibit differential distributions across high-temperature extreme reefs versus more typical classic reefs (Figure 1). These lineages exhibit differences in their strain-level photobiont associations and bacterial community compositions. Lineages also differ in terms of optical traits, with those predominantly found in extreme reefs having lower light absorption efficiency (Figure 6). Together, these factors interacted to shape thermal tolerance; lineages typically observed on extreme reefs exhibited reduced bleaching and mortality during thermal challenge. Interestingly, thermal tolerance also differed between lineages inhabiting extreme reefs, suggesting that host identity is a more important driver of thermal tolerance than reef type (Figure 7). Together, these findings show that co-occurring cryptic coral lineages, although visually indistinguishable, can exhibit strong functional variation with important implications for conservation and management under future climate change.

### Some cryptic lineages may represent widespread species

The three identified cryptic lineages (L1-L3) of massive *Porites* differed in their relative abundances among classic and extreme reefs (Figure 1). Based on these distributions, we posit that L1 is a “classic” reef specialist, L2 is a generalist lineage, and L3 is an “extreme” reef specialist. Our sampling efforts support the findings of Rivera et al. (2022) that lineages of massive *Porites* differ in their distributions among Palau’s Rock Island reefs. We infer that L1 is equivalent to the dark blue (DB) lineage in Rivera et al. (2022); L2 is equivalent to light blue (LB); and L3 is equivalent to red (RD). A fourth lineage (pink (PI)), reported by Rivera (2022) to be more abundant at sites outside of the Rock Islands, was not represented in our study. These data, combined with previous reports from across the Pacific Ocean, show that lineages specialized to nearshore and offshore habitats (Afiq-Rosli et al., 2021; Boulay et al., 2014; Schweinsberg et al., 2016; Tisthammer et al., 2020), as well as different depths (Voolstra et al., 2023) are a common feature on Pacific reefs.

Combined analysis of two 2b-RAD satasets showed that two of the Palauan lineages (L1, L2) are more closely related to populations ∼4300 km away in Kiritimati, than to co-occurring lineages on the same reefs (Figure S5). Both locales harbored lineages that were highly diverged from all sampled populations, suggesting that while some cryptic *Porites* lineages may have wide distributions (*i.e.,* L1, L2 in Palau), others may be locally endemic (*i.e.,* L3 in Palau). The levels of genetic differentiation observed between Palauan and Kiritimatian populations may be a common pattern of isolation by distance in gonochoric broadcasting species (Aichelman & Barshis, 2020; Nunes et al., 2011). Hence, we posit that L1 and *Pkir*-*2,* as well as L2 and *Pkir-1*, likely represent distinct populations of two lineages with limited gene flow. Based on morphological data (Figure 2, 8), the L1-*Pkir-2* lineage is likely most related to *P. australiensis*, and L2-*Pkir-1* to *P. lobata.* L3 appears to be an additional species that exhibits morphological similarities to L2 and *P. lobata*. Of note, while hybrid colonies were not found in this survey or in the recent work in Kiritimati (Starko et al., 2023), Rivera et al (2022) did identify three adult colonies that were inter-lineage hybrids, suggesting that these lineages can hybridize to some extent, albeit rarely.

Of interest, while L3 was the most genetically distant lineage, it was morphologically indistinguishable from L2 and challenging to distinguish from L1. This signals a decoupling of morphological and genetic differentiation, in line with previous surveys showing that samples matching the morphologies of *P. lobata*, *P. lutea*, *P. annae*, *P. solida*, and *P. harrisoni* together form a single clade in which morphotypes are genetically indistinguishable (Terraneo et al., 2021). Moreover, across Singaporean reefs, phylogenetic analyses of *Porites* could not resolve morphologically identified species, forming a complex that included *P*. *australiensis*, *P*. *lobata* and *P*. *lutea* (Quek et al., 2023; Quek & Huang, 2019). Further highlighting the taxonomic incongruencies among massive *Porites*, colonies matching the morphology of *P. lutea* were identified in three distinct genetic clades of *Porites* across the Pacific, and two in *P. lobata* (Forsman et al., 2009). These phylogenomic analyses indicate that massive *Porites* consists of diverse species complexes, potentially each with plastic corallite morphologies. Yet, our findings suggest that morphological characteristics can, in some cases, still be useful for distinguishing genetic lineages. Broad-scale genetic sampling of morphologically similar colonies across various habitat types and depths coupled with skeletal morphology assessments by trained taxonomists is necessary to fully disentangle the taxonomy and distributions in Pacific massive *Porites*, as well as other coral genera.

### Locally specialized and lineage-specific photobiont associations

Associations with distinct genera, species, or even strains of photobionts can have tremendous impacts on coral holobiont function and physiology (LaJeunesse et al., 2018; Thornhill et al., 2017). Yet, the potential for association with diverse photobionts may be limited among species of massive *Porites* given that they transmit photobionts maternally (Bennett et al., in review; Forsman et al., 2020). Nonetheless, some corals employing parental photobiont provisioning can also acquire photobionts via horizontal transmission (Byler et al., 2013; Quigley et al., 2018; M. J. H. Van Oppen, 2004). We found that all three *Porites* lineages in Palau harbored *Cladocopium* C15 photobionts, yet strain-level variation was evident amongst lineages, as well as between classic and extreme sites. These findings indicate population structure in the photobiont and suggest that some lineages may also engage in horizontal transmission of photobionts, which could facilitate environmental acclimation (Figure 3, 8).

Given that L1 colonies harbored a distinct DIV at extreme reefs not found in colonies of the same lineage on classic reefs (Figure 3), as well as background abundances of *Symbiodinium*, we infer that L1 is a host-photobiont generalist, associating with locally available photobiont strains across environments. By comparison, L2 was more promiscuous in its associations and no clear patterns were observed across reef types. Most L3 colonies (N=8/10) associated with two DIVs that were not found in the other lineages. This pattern is likely the result of vertical transmission (Forsman et al., 2020; Scott et al., 2024), supporting host-photobiont specialization. Similar variation in host-photobiont fidelity at the strain level was identified among cryptic *Porites* lineages in Kiritimati (Starko et al., 2023), where one lineage hosted a distinct strain of C15 compared to the other lineages. However, a marine heatwave eroded this tight partnership, showing that abrupt environmental change can alter these associations (Starko et al., 2023).

Our results also suggest that some photobiont DIVs may be beneficial for survival of massive *Porites* lineages on extreme reefs. For instance, one photobiont DIV (C15.C93a) was common at extreme sites among colonies of all lineages (L1, *n* = 4/4; L2, *n* = 12/22; L3 *n* = 2/10), potentially indicating selection of locally beneficial host-photobiont pairings. Preliminary analysis of the effect of photobiont DIV on colony thermal tolerance in the thermal challenge experiment suggests that colonies with this strain maintained higher Fv/Fm than colonies hosting other strains (Figure S11). However, additional work is needed to confirm this hypothesis. Alternatively, the observed patterns may simply indicate limited photobiont dispersal between extreme and classic sites (Golbuu et al., 2012).

### Microbiome regulation strategies differ among lineages

Bacterial assemblages fulfill important services in coral holobionts, and association with locally beneficial bacteria can help corals respond to changing environments (Peixoto et al., 2021; Voolstra et al., 2024). Yet, corals differ in their potential to switch or shift between bacterial microbiome members, *i.e.,* microbiome flexibility, affecting their potential for acclimatization (Ziegler et al., 2019). We found that bacterial communities differed between cryptic lineages and reef types (Figure 4, 5). Patterns of bacterial community assembly also differed between reef types in lineage-specific ways, suggesting differences in microbiome flexibility. For instance, L1 microbiomes were more variable at extreme sites than at classic sites whereas L2 colonies maintained relatively stable microbiomes regardless of reef type. Based on these findings, we posit that L1 is a microbiome conformer and L2 is a microbiome regulator (Ziegler et al., 2019). Microbiomes associated with L3 appeared highly variable and diverse, but given its limited distribution, it is not possible to disentangle whether this is a general feature of this lineage or the results of local stochasticity.

We observed a loss of Endozoicomonadacae in L1 at extreme sites, whereas L2 maintained relatively high abundances of this bacterial family (Figure 5). Notably, none of the sampled L3 colonies hosted appreciable proportions of Endozoicomonadaceae. A loss of this bacterial family has been documented in corals living in extreme environments and was attributed to environmental stress (Camp et al., 2020; Pantos et al., 2015), but has also been observed in corals regularly fed while living in captivity, potentially indicating reduced need for this putative nutritional symbiont (Barreto et al., 2021; Pogoreutz & Ziegler, 2024). While paucity of Endozoicomonadaceae in the microbiomes of L1 and L3 may thus indicate microbiome disruption due to environmental stress, it could also indicate reduced need for this partner due to higher heterotrophic feeding in nearshore habitats with increased suspended particles (Anthony & Fabricius, 2000; van Woesik et al., 2012). Future work investigating links between heterotrophy and Endozoicomonadaceae abundances among these lineages is warranted.

Individual Endozoicomonadaceae ASVs exhibited surprisingly heterogeneous distributions among lineages and reef types (Figure 5B). Some ASVs were more strongly associated with certain lineages (*e.g.,* ASVs 10, 12, and 20), potentially indicating host-specialization. Comparable patterns of host-lineage specialization were observed among Endozoicomonadaceae strains associated with genetically distinct populations of *Stylophora pistillata* at various locations (Buitrago-López et al., 2023; Neave et al., 2017). Some ASVs appeared to be more widely distributed and may therefore be cosmopolitan partners of massive *Porites* (ASV 1). Such widespread bacterial associations appear commonplace in corals that employ horizontal transmission of microbial communities and may indicate that some abundant Endozoicomonadaceae ASVs are taken up from the environment (Neave et al., 2017). Other strains that were more abundant on one reef type (*e,g.,* ASV6 on classic sites) may be more closely tied to the local environment than host lineage (*i.e.,* classic sites in this study).

### Optical adaptations of holobiont lineages to distinct reef types

Corals receive most of their nutrition from their photobionts (Muscatine, 1990). As a result, they have evolved mechanisms to optimize light harvesting in their environments (Enríquez et al., 2005, 2017; Terán et al., 2010). A common feature of cryptic coral lineages is that they are separated across habitats that differ in terms of light intensities, such as across depths or distance from shore (Grupstra et al., 2024), suggesting that they may be adapted to optimize local light conditions. Although we did not observe differences in light intensity between classic and extreme reefs over a 16-day measurement period in 2022 (Figure S2), previous work has shown that Palau’s extreme reefs have higher light attenuation due to increased densities of suspended particles (van Woesik et al., 2012). Low light levels on Palau’s extreme reefs have also been hypothesized to mitigate stress caused by high temperatures and result in the selection of low light-adapted coral species (van Woesik et al., 2012).

We found that lineages differed in terms of key traits that are correlated with light harvesting strategies that can inform resistance to temperature and light stress. For instance, polyp density is associated with light scattering potential of the coral skeleton (Figure 6A; Gómez-Campo et al., 2024). Importantly, lineages differed in terms of Chl*a* densities and light absorption efficiency (Figure 6, 8). Low Chl*a* concentrations coupled with high light absorption efficiency, as seen in L1, results in a reduction in self-shading of the photobiont cells and increased light scattering of the coral skeleton, which is commonly associated with high-light coral phenotypes and leads to higher energy production (Enríquez et al., 2005; Gómez-Campo et al., 2022; 2024). However, a reduction in self-shading also increases light stress in photobionts, increasing bleaching probability in high temperatures. This combination of factors likely underpins why L1 is less abundant on extreme reefs where high temperatures increase photobiont stress, raising their probability of bleaching and mortality.

**Figure 8.**
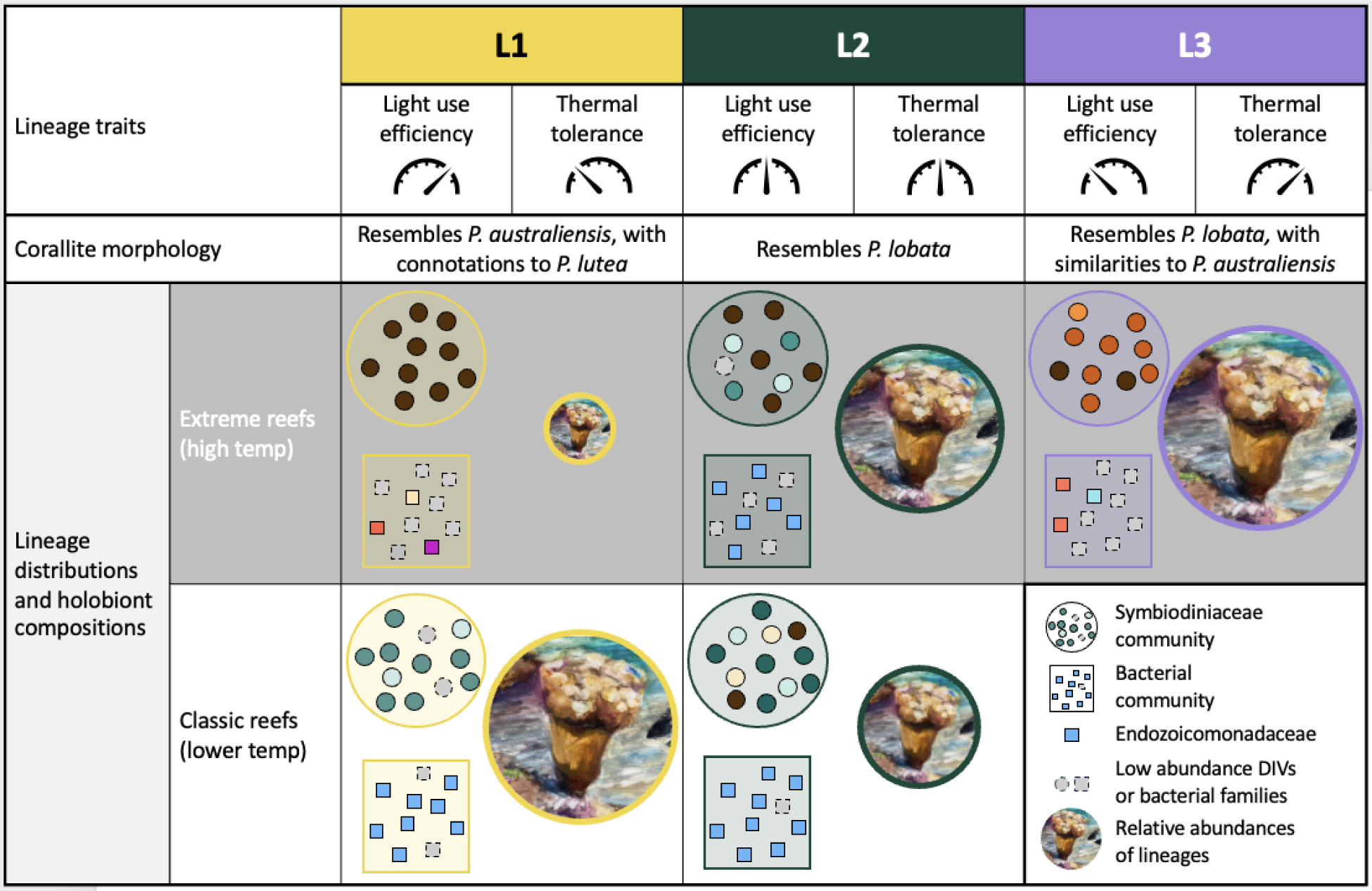
Overview of lineage traits, morphologies, distributions, and holobiont compositions. Lineages of massive *Porites* differ in their light use efficiency and thermal tolerance (low (pointing left), medium (pointing up) and high (pointing right) values). Analyses of corallite traits indicate subtle morphological differences among lineages. Lineages also differ in terms of their abundances and photobiont (Symbiodiniaceae, circles) and bacterial (squares) associations across extreme and classic reefs. Coral icon sizes are scaled per lineage based on frequencies from Figure 1. Gray circles and squares represent low abundance photobiont DIVs or Bacterial families, respectively. Colors of circles and squares roughly correspond to colors in figures 3 and 5. Note that L3 corals were never observed on classic reefs. Coral reef artwork by Kimberly Collins Jermain.

Low-light phenotypes, on the other hand, are generally characterized by higher Chl*a* concentrations, which increases the likelihood of photon capture, but simultaneously results in higher chlorophyll self-shading, lower light absorption efficiency, and reduced total photosynthesis (Enríquez et al., 2005; Gómez-Campo et al., 2022; Gómez-Campo et al., 2024; Mass et al., 2007; Winters et al., 2009). L3 colonies match these traits, and this likely explains why this lineage is restricted to extreme reefs: high Chl*a* concentrations can be detrimental in high-light environments through increased production of reactive oxygen species (ROS) that damage photobiont and host cells (Brown, 1997; Cruz De Carvalho, 2008; Weis, 2008). Reduced photosynthesis may also partially explain why L3 colonies have lower growth rates than colonies from co-occurring lineages (Rivera et al., 2022).

L2 colonies had intermediate levels of Chl*a* and light absorption efficiency compared to L1 and L3, promoting survival in diverse habitats and supporting the hypothesis that this lineage is a habitat generalist. Of note, a lack of differences in Chl*a* concentrations and light absorption efficiency among L2 colonies sampled in extreme and classic reefs (Figure 6) suggests that these traits may not be plastic, limiting the ability of massive *Porites* lineages to adjust to environmental change, and potentially resulting in the elimination of lineages not suitable for survival under future climate change conditions.

### Response variation among holobiont lineages to thermal challenge

Identifying adaptations and acclimatory mechanisms that shape responses to thermal challenge is critical to predicting the effects of rising ocean temperatures on coral communities and can aid restoration efforts (Barshis et al., 2018; Thomas et al., 2018). This is especially important for cryptic coral lineages that were long assumed to be functionally similar (Grupstra et al., 2024). We found that distributions, holobiont compositions, and light absorption efficiency correlated with thermal tolerance in cryptic coral lineages, as tested using a common garden thermal challenge (Figures 3-8). These lineage-specific thermotolerances are consistent with Rivera et al. (2022), who reported more stress bands during a bleaching event in 1988 in L1 (DB, 68%), relative to L2 (LB, 22%) and L3 (RD, 25%). Together, our findings suggest that extreme reefs select for locally beneficial holobionts with adaptive light harvesting traits that help corals delay bleaching under thermal challenge (Gómez-Campo et al., 2022; Scheufen, Iglesias-Prieto, et al., 2017; Scheufen, Krämer, et al., 2017; Swain et al., 2016, 2018).

We also found critical differences between L2 and L3 in terms of mortality and Fv/Fm under thermal challenge, showcasing important response variation and distinct modes of holobiont adaptation among lineages from the same environment. It is likely that the factors that restrict L3 to extreme reefs, such as high fidelity for specific photobiont strains and increased specialization to low-light habitats (Figures 3-8), support its survivorship under thermal stress (Howells et al., 2012, 2020; Swain et al., 2016). On the other hand, association with more diverse photobiont strains, along with intermediate light harvesting efficiency, helps L2 live on diverse reef types, but this generalist strategy likely reduces survivorship of this lineage under thermal challenge. These findings also suggest that future marine heatwaves are likely to favor the survival of L3 compared to L2, warranting further investigation into the molecular mechanisms underpinning thermal tolerance in this holobiont, and establishing this lineage as an important resource for conservation and restoration efforts focused on reefs with high turbidity.

One important question remaining is whether there are tradeoffs to the increased thermal tolerance in lineages of massive *Porites*. For example, colonies of massive *Porites* from extreme mangrove habitat had lower gene expression variation, reduced skeletal density, and increased porosity compared to populations on “classic” reefs (Scucchia et al., 2023). In Palau, Rivera et al., (2022) found that L3 had lower skeletal density, calcification, and extension rates than the other lineages. Our findings also suggest that L3 may be less well-equipped to handle light stress. Identifying such trade-offs in lineages with increased thermal tolerance will reveal important considerations for restoration efforts (but see Lachs et al., 2023).

It is also important to note the experimental design limitations in this study. The thermal challenge experiment limits our ability to disentangle lineage and environment, given that all L1 colonies were collected from classic reefs whereas L2 and L3 colonies were collected from extreme reefs. We also did not control or measure light intensity during the experiment. Future experiments testing acclimatization to extreme reefs and host/photobiont genetics, for example using reciprocal transplant experiments, will help disentangle these effects.

## Conclusions

Extreme reefs can offer a glimpse into the potential future for corals, revealing adaptations and microbial partnerships that can aid survival under future climate change conditions. Our findings suggest that extreme reefs with high temperatures and high light attenuation promote associations with locally beneficial partners as well as the evolution and proliferation of optical traits that reduce light stress. Importantly, lineages appeared to employ distinct adaptations and acclimatory mechanisms to survive on extreme reefs, showing that there is no “one size fits all” mechanism that promotes survival. Some of the thermotolerant cryptic lineages discussed here, as well as in other recent works, may be suitable candidates for restoration efforts aimed to increase the abundances of thermally tolerant corals on reefs threatened by climate change. The use of slow-growing stress-tolerant species, such as massive *Porites* in particular, represent underutilized resources that can complement current restoration efforts (Guest et al., 2023).

Altogether, our findings emphasize the importance of resolving host genetic variation using molecular methods when conducting ecological experiments or determining “winners’’ and “losers’’ following bleaching events (Loya et al., 2001). L2 and L3 colonies are micromorphologically indistinguishable. Yet, response variation to thermal stress events among these lineages may result in heterogeneous mortality. Such differences in mortality among lineages of massive *Porites* were recently reported in Kiritimati where one lineage experienced 75% mortality while overall mortality of the other lineages was only 20% (Starko et al., 2023). Our findings show that assemblages of *Porites* in the Pacific are composed of combinations of wide-spread and distinct local, potentially endemic, lineages. This suggests that heterogeneous responses to marine heatwaves are likely commonplace, potentially resulting in the elimination of yet unknown lineages. Combined with recent work showing comparable differences in thermal tolerance between cryptic lineages across the anthozoan tree of life (Grupstra et al., 2024), these findings demonstrate that cryptic diversity is a key factor driving patterns of holobiont structuring as well as heterogeneous responses to ocean warming. Identifying and accounting for cryptic lineages is of key importance when quantifying phenotypic variation and planning restoration strategies.

## Acknowledgements

We would like to thank MJ Shanks for assistance with fieldwork and PICRC staff, including Joy Schmull-Sam, Louw Claassens, Arius Merep, and Rodney Kazuma for logistical support. Special thanks go out to Kimberly Collins Jermain for leading outreach activities with Palauan students and painting the artwork used in Figure 8. We also thank Justin Scace and Joel Sparks for support with photography of coral skeletons and SEM, respectively. Members of the Coralassist lab are thanked for valuable discussions and insight. Davies lab members provided valuable feedback on the conducted analyses and figures.

## Author contributions

Study design – CGBG, SWD, KMK, MJB, MA; Data collection – CGBG, KMK, MJB, KMK, MA, DJJ, JPDA, KJGC, IMR; Sample processing - CGBG, KMK, AKH, AMH; Data analysis – CGBG, JEF, HEA, DJJ, SWD, KMK; Manuscript writing – CGBG, DJJ, JPDA; Manuscript editing – all authors.

## Data availability

Raw host-targeted (2b-RAD) sequencing data were uploaded to the sequence read archive (SRA) under accession xxxx. Raw microbiome-targeted sequencing data (combined ITS-2 and 16S rRNA gene amplicon sequencing) were uploaded to the SRA under accession xxx. All other datafiles are included as supplementary datafiles. All code to replicate the analyses conducted here has been uploaded to GitHub: https://github.com/xxxxx

## Permits

Sample collection was authorized by scientific research permits issued by the Palau Ministry of Natural Resources, Environment, and Tourism (RE-21-17, RE-22-17, RE-22-24, RE-23-09) and Koror State Government (69, 72, 78, and 83). Importation of coral samples to the United States was conducted with authorization from the Palau Bureau of Marine Resources and U.S. Fish and Wildlife Service (CITES permits PW22-042, PW22-162, PW23-089, PW23-090).

