## Supplementary for "Distinct modes of holobiont specialization among cryptic coral lineages"

### Running head:

Specialization in cryptic coral lineages

### Authors:

Carsten G.B. Grupstra^1^*, Kirstin S. Meyer-Kaiser^2^, Matthew-James Bennett^3^, Maikani O. Andres^4^, David J. Juszkiewicz^5^, James E. Fifer^1,6^, Jeric P. Da-Anoy^1^, Kelly J. Gomez Campo, Isabel Martinez Rugerio, Hannah E. Aichelman^1^, Alexa K. Huzar^1^, Annabel M. Hughes^1,7^, Hanny E. Rivera^1^, Sarah W. Davies^1^*

*corresponding authors

*^1^Department of Biology, Boston University, Boston, MA, USA.*

*^2^Biology Department, Woods Hole Oceanographic Institution, Woods Hole, MA, USA*

*^3^MARE, Guia Marine Laboratory, Faculty of Sciences, University of Lisbon, Cascais 2750-374, Portugal*

*^4^Palau International Coral Reef Center, Koror, Palau 96940*

*^5^Coral Conservation and Research Group (CORE), Trace and Environmental DNA Laboratory (TrEnD), School of Molecular and Life Sciences, Curtin University, Bentley, Western Australia 6102, Australia*

*^6^Department of Ecology, Evolution, and Marine Biology, University of California San Diego*

*^7^Northeastern University Marine Science Center, Nahant, MA, USA*

### Supplementary Materials

#### Supplementary Methods

##### Site selection and sample collection

Water temperature and light levels were recorded every 30 minutes at each site using Hobo Tidbit and Pendant loggers (Onset, Wareham, USA). Temperature loggers were deployed from November 2021 to May 2022, but light loggers were only deployed for 16 days (during sampling trips) to prevent biased data acquisition from biofouling.

##### DNA extraction, library prep, and sequencing

Tissue samples from all coral colonies (sampled in November 2021 and April 2022) were crushed with a sterile razor blade, and DNeasy Blood and Tissue kits (Qiagen) were used to isolate DNA from the resulting homogenate according to the manufacturer’s instructions, with one modification: the lysis step was conducted overnight. Isolated DNA was then cleaned with a Zymo Clean and Concentrator kit (Zymo Research, CA). DNA was quantified using a Qubit fluorometer (Invitrogen), standardized to 25 ng μL^-1^, prepared for 2b-RAD sequencing according to (Wang et al., 2012), and sequenced across one lane of Illumina HiSeq 2500 using single-end 50 bp sequencing at the Tufts University Core Facility (TUCF) Genomics. Five technical replicates were included in the library preparation to aid the downstream identification of clonemates. For samples collected in April 2022, we used the same approach with modifications for fewer loci (reduced representation) (Wang et al., 2012).

##### 2b-RAD data analysis

For samples collected in November 2021, raw reads were deduplicated and trimmed using the FASTX toolkit (http://hannonlab.cshl.edu/fastx_toolkit). Reads under 25 bp in length or with quality scores <15 were discarded. Following Rippe et al. (2021), photobiont reads were removed by discarding reads that mapped to concatenated Symbiodiniaceae genomes (*Symbiodinium* (Aranda et al., 2016)*, Breviolum* (Shoguchi et al., 2013)*, Cladocopium* (Dougan, 2020), and *Durusdinium* (Dougan, 2020) with Bowtie2 v2.4.2 (Langmead & Salzberg, 2012). The remaining host reads were then mapped to the *Porites lobata* genome (Noel et al., 2023). Genotyping was performed using ANGSD v0.923 (Korneliussen et al., 2014). Filters that were used across all analyses included loci that were present in ≥ 80% of individuals, and a minimum read depth of 6 across all samples. Triallelic sites were removed. Reads had a minimum quality of 25, minimum mapping quality of 20, with a strand bias p-value of 1e-5 and a heterozygosity bias p-value of 1e-5. Clones were detected using hierarchical clustering based on pairwise identity by state (IBS) with an additional minor allele frequency (MAF) filter of 0.05. Technical replicates providedthe clone detection threshold, and one member of each clone pair was removed for downstream analyses.

After the first 2b-RAD run, a total of 75 samples remained after quality control and technical replicate removal. These libraries were selected for further population genomic analyses due to a higher proportion of the genome covered compared to reduced RAD samples. For all population genomic analyses an additional MAF filter of 0.05 was added, with the exception of site frequency spectrum (SFS) based analyses. Admixture was estimated using NGSadmix; admixture plots were then created using a custom R script (<https://github.com/z0on/2bRAD_denovo/blob/master/admixturePlotting_v5.R>). Principal Component Analysis (PCA) was conducted using a covariance matrix based on single-read resampling calculated in ANGSD. Admixture results were visualized using the K with the least cross validation error reported from ADMIXTURE. These analyses demonstrated the presence of three distinct lineages amongst our six sampling sites. F_ST_ was estimated between these lineages using ANGSD before and after outlier loci were removed using Bayescan (Foll & Gaggiotti, 2008). To determine the extent to which the lineages of *Porites* massive in Palau are related to recently discovered cryptic lineages of massive *Porites* in Kiritimati (located ~4300km eastward from Palau), we re-analyzed our 2b-RAD data along with 2b-RAD data published in Starko et al., (2023). Global F_ST_ was calculated from each lineage's SFS generated by first calculating the site allele frequency (SAF) in ANGSD with no MAF filter and then unfolded (using the *Porites lobata* genome (Noel et al., 2023) as an ancestral reference) SFS in winsfs (Rasmussen et al., 1982). Weighted F_ST_ was then calculated using ANGSD based on 13,443 loci. Outliers were not removed for this analysis because the dataset from Kiritimati had fewer loci.

Samples collected in Palau in April 2022 were sequenced in a later 2b-RAD run with a reduced representation design. We also included 15 samples from 2021 that failed initial 2b-RAD sequencing. 15 samples from 2021 with known lineage were resequenced (*n* = 5 per lineage) to facilitate lineage assignments. Lineage assignments were made using hierarchical clustering based on pairwise IBS. Data from this run were not included in the population genetic analyses (*e.g.,* F_ST_ and admixture estimation analyses) presented here given the differences in sequencing efforts, but lineage assignments from this run were used to inform lineage distributions and downstream microbial community analyses and thermal challenge experiment results. A Chi-squared test was used to test the hypothesis that the three identified lineages were unevenly distributed between extreme and classic sites in Palau using data from both sequencing runs.

##### Micro-morphological observations

Micro-morphological characterisation was conducted on 19 skeletal fragments (2 x 2 cm^2^) of *Porites*, each originating from distinct colonies (classic *n =* 8; extreme *n =* 11). Specimens were imaged with a Canon EOS 60D camera with a macro lens (MP-E 65 mm) mounted on a camera rail (COPYMATE II, Bencher). Photos were focus-stacked using HeliconFocus v. 8.2.10. A subset of samples (L1, *n =* 6; L2, *n =* 4; L3, *n =* 3) was also imaged using a Phenom proX scanning electron microscope (SEM). Images were exchanged in a blinded batch file for morphological examination, devoid of lineage information to limit confirmation bias. Macro-morphological observations focused on key diagnostic features pertaining to the calice, wall, septa, and columella (Bernard, 1905; Budd et al., 1994; Vaughan, 1918; Veron & Pichon, 1982). Species-level identification was determined by observations of the diagnostic features and referring to published morphological descriptions of *Porites* in the Indo-Pacific, with a focus on species recorded in Palau, Micronesia (Crossland, 1952; Eguchi, 1938; Vaughan, 1907, 1918; Veron, 1985, 1986; Veron & Pichon, 1982; Wells, 1954). Post-examination, species level identifications were then compared to the *a priori* lineages (L1, L2, L3) defined by molecular analyses.

##### Microbial community sequencing

Photobiont communities were characterized in samples collected in November 2021 through sequencing of the internal transcribed spacer region 2 (ITS2) region using *SYM_VAR_5.8S2* and *SYM_VAR_REV* primers (Hume et al., 2015, 2018). The PCR profile included 26 cycles of 95℃ for 40 s, 59℃ for 2 min, 72℃ for 1 min and a final extension of 72℃ for 7 min. A negative control was included in the initial amplification but failed to amplify, so it was not included in downstream library preparations. Successful amplifications were cleaned using the GeneJET PCR Purification kit (ThermoFisher Scientific) and a second PCR was conducted to attach Illumina MiSeq dual barcodes to the PCR product before samples were pooled. Volumes for pooling were based on the visualization of barcoded sample band intensity on a 1% agarose gel. This pool was cleaned using the GeneJET PCR Purification kit, gel extracted, and submitted for sequencing as described below.

To characterize bacterial communities, the V4 region of the 16S rRNA gene was amplified from the same samples via PCR using Hyb515f (Parada et al., 2016) and Hyb806R (Apprill et al., 2015) primers and the following PCR profile: 35 cycles of 95 ℃ for 40 s, 65 ℃ for 2 min, 72 ℃ for 1 min and a final extension of 7 min. Subsequent PCR amplification, cleaning, dual-barcoding, and gel extraction followed the same protocol described for ITS2 with the inclusion of three negative controls, which were also submitted for sequencing. ITS2 and 16S pools were quantified and combined in a 1:3 ratio, respectively. Libraries were sequenced together on Illumina MiSeq (paired-end 250 bp) at Tufts University Core Facility (TUCF) Genomics.

##### Microbial bioinformatics processing and analysis

Sequences with adaptor contamination were removed and raw 16S and ITS-2 sequences were separated based on primer sequences using bbduk following Bove et al. (2023). Raw ITS-2 reads were processed by Symportal (Hume et al., 2019) to produce defining intragenomic sequence variant (DIV) profiles for each coral colony. Two samples with <1,000 reads were removed, as well as one outlier sample with >1 million reads; the remaining samples (*n =* 73) had an average of ~5,500 reads per sample (min: 1,116; max: 16,931). All samples were dominated by one of ten *Cladocopium* C15 types, and four samples hosted low abundances of *Symbiodinium* A3 sequences. To test for differences in dominant C15 DIV associations among lineages and reef types, we fitted cumulative link models (clm) using the package ordinal (v2022.11-16) on a dataset from which background abundances of A3 were removed. We included lineage and DIV as fixed effects with an interaction, and site was included as an additional fixed effect to determine site-level variation (final model: DIV~Lineage*Reef Type + Site).

Primer sequences were removed from raw 16S reads using cutadapt 4.1 (Martin, 2011). Quality filtering, denoising, merging, and taxonomy assignments were conducted with DADA2 against the *Silva* v. 138.1 database (Quast et al., 2012). Only merged reads of lengths 251-256 bp were maintained, and chimeras were removed using the “consensus” method. Mitochondrial and chloroplast reads were removed, and bacterial reads were filtered with the subset_taxa command in phyloseq v1.38.0 (McMurdie & Holmes, 2013). Contaminant reads were removed using the prevalence method in the decontam package v1.14.0 (Davis et al., 2018), leaving a total of *n =* 48 (out of 90) individuals for downstream analyses. Diversity metrics (Shannon, Simpson) were calculated based on the non-rarefied dataset using the estimate_richness function. We tested for differences between lineages and sites using a linear model with an interaction between lineage and reef type. Including site as a random effect (using lmer) resulted in a model convergence failure; it was therefore removed from the final model.

Bray-Curtis distances were calculated based on data from which rare (<10 reads) ASVs were removed (2361 taxa, *n =* 48). Unrarefied data were total-sum-normalized as advised by (McKnight et al., 2019) and Bray-Curtis distances were calculated and used for visualization in a non-metric multidimensional scaling (nMDS) plot using the distance and ordinate (K = 2) functions in phyloseq. We tested for differences in bacterial community composition among lineages and reef types while including collection site as an effect nested within reef type using PERMANOVA with the function adonis2. We tested for differences in dispersion among sampling groups (five groups: L1 classic, L1 extreme, L2 classic, L2 extreme, L3 extreme) using betadisper because PERMANOVA is sensitive to differences in dispersion between sampling groups if replication is uneven among groups (Anderson & Walsh, 2013). Lastly, the top 10 most abundant Endozoicomonadaceae ASVs were filtered and associations between lineages and specific Endozoicomonadaceae strains were explored using a bubble plot.

##### Analysis of polyp densities and colony coloration

To identify potential morphological differences that could inform ecological function, we quantified polyp densities in a subset of coral colonies that were recovered after initial tagging (L1 classic *n =* 10, extreme *n =* 4; L2 classic *n =* 3, extreme *n =* 10; L3 classic *n =* 0, extreme *n =* 6). Photographs were taken using an Olympus TG-6 camera set to macro mode under natural lighting with a size and color standard to calibrate coloration data. Polyp densities were quantified by counting the number of polyps in 4 mm^2^ sections in ImageJ v1.53K. We tested for differences in polyp densities among lineages using a linear model with an interaction between reef type and lineage. Pairwise comparisons among lineages (i.e., regardless of reef type) were conducted using emmeans on a model without the interaction. To test for differences in polyp densities between reef types within lineages, we filtered data for just L1 or L2 and used a linear model with reef type included as a fixed effect.

##### Measurement of lineage-specific light harvesting properties

In May 2023, 20 tagged colonies of known lineage and reef type (classic: L1 *n =* 5, L2 *n =* 4, L3 *n =* 0; extreme: L1 *n =* 0, L2 *n =* 6, L3 *n =* 5) were transported to Boston University and fragmented. Fragments were then acclimated for 63 days in aquariums (28˚C, 33-35 ppt salinity, 10 hr dark: 14 hr light cycle, 100-115 PAR) before transportation to Pennsylvania State University.

Live samples were placed in a black container filled with filtered seawater. The container was illuminated with uniform diffuse light emitted from a semi-sphere coated with barium oxide (BaO) and positioned above the sample and the black container. Inside the black container, a submersible LED ring was placed around the coral, and additional halogen lamps and violet-blue LEDs were used to enhance the illumination in the red-infrared and violet-blue regions, respectively, which was then reflected by the semi-sphere. To collect the reflected light, a 2 mm diameter fiber-optic probe was placed at a 45° angle and 1 cm away from the coral surface. Measurements were conducted between 400 and 750 nm using a Miniature Ocean Optics USB 4000 spectroradiometer (Ocean Optics Ltd, FL). Calibration was performed using the reflectance of a bleached coral skeleton of the corresponding species. To determine the optical properties of coral, we measured coral reflectance (R)  and absorptance (A = 1 - R) following the methods described by Enríquez et al. (2005) and Vásquez-Elizondo et al. (2017).

The specific absorption coefficient normalized to chlorophyll a at 675nm (a*_Chl_*_a_* , m^2^ Chla ^−1^) was calculated according to Enríquez et al. (2005), using the equation:

𝑎∗ = (𝐷𝑒/𝜌)⋅𝑙𝑛(10)

where De is the estimated absorbance value, calculated as De = log(1/R), and ρ the chlorophyll a content per projected area in mg Chla m−2.

To quantify Chl*a* concentrations,  coral samples were airbrushed with filtered seawater (0.45µm), and subsequently, the coral tissue slurries were homogenized using a tissue homogenizer (T 10 basic Ultra-Turrax, IKA). Photobiont cells were concentrated through centrifugation at 2000 rpm for 5 minutes. Afterwards, 10ml of supernatant was collected and stored at −20°C for protein determinations. The remaining pellet was resuspended in filtered seawater and utilized for chlorophyll *a* density assessments. Pigment extraction was carried out with acetone/dimethyl sulfoxide (95:5, vol/vol). The samples were kept in darkness at 4°C for 24 hours and then centrifuged before spectrophotometric determinations were made. Chlorophyll determinations were measured using a Miniature Ocean Optics USB4000 (Ocean Optics Ltd, Fl) and a fixed optical geometry. The equations provided by Jeffrey & Humphrey (1975) for dinoflagellates were used to calculate the final chlorophyll content. Pigment contents were standardized by coral surface area (McLachlan & Grottoli, 2021a).

We tested for differences in chlorophyll concentration and a*_chla_ values among lineages using a linear model with reef type and lineage as fixed effects. An interaction between reef type and lineage was not included because we had samples from both reef types for just one lineage (L2). Pairwise comparisons among lineages were conducted using emmeans with a Bonferroni p-value correction. Model assumptions were verified visually.

##### Thermal challenge experiment

A 25-day common garden heat challenge experiment was conducted to test for differences in thermal tolerance between the three *Porites* lineages (see Supplementary Methods)*.* Briefly, 24 tagged colonies of known lineage were included from five study sites (Table S1). L1 colonies were collected from classic sites and L2 and L3 colonies were collected from extreme sites (Table S1). We initially aimed to include L1 and L2 colonies from both extreme and classic sites, but were unsuccessful. Two cores (~5.7 cm diameter) were extracted from each colony using a hole saw, and cores were haphazardly assigned to one of three heat challenge treatment tanks or one of three control tanks (6 tanks total, containing ~30 L unfiltered seawater each) such that one core from each colony was represented in each treatment. 50% water changes were conducted daily. Temperature was measured every 30 minutes using temperature loggers (Onset, MA). Mean temperatures in control tanks were maintained at 29.5 °C±0.1 °C. Temperatures in the heat challenge treatment tanks were ramped by ~3°C over seven days, followed by a hold at 32.4±0.2°C for 12 days. On day 19, temperatures were increased by an additional ~1°C (to 33.5±0.1 °C) to simulate an extreme thermal stress event, which was maintained until day 25.

Coral cores were declared dead when no visible tissue remained, no typical fluorescence under blue light was observed, and cores were visibly covered with biofilm or turf algae. Maximum PSII photochemical efficiency (Fv/Fm) was measured daily between 20h00 and 21h00 (after at least 1.5 hours of darkness) using a Junior PAM fluorometer (Walz, Germany) for the first six days of the experiment and then every second day thereafter. Additional Fv/Fm measurements were collected on day 19 and day 21. To quantify rates of tissue paling, all fragments were photographed with a color standard prior to start of the experiment (day 0), as well as every ~5 days at ~14h00 on each day to standardize lighting conditions over the duration of the experiment. Bleaching was quantified as the intensity in the gray channel using the Fiji software package (sensu McLachlan & Grottoli, 2021b).

#### Supplementary Results

##### Limited corallite diagnostic features are discernable among lineages

#### L1

L1 corallite morphology is characterized by shallow calices with gentle sloping septal margins (Figures 2B, C, D). The walls, typically straight and of medium to thick composition, form a ridge with rough granulated mural denticles (Figure 2B, C). The septa appear thick, and the interseptal loculi tend to be narrower or equal to the septa (Figure 2B, C, D). Occasionally, the outer ends of the septa bifurcate, corresponding to a mural denticle (Figure 2C). The inner synapticular ring is mostly complete, aligning with the position of the pali. Typically, there are eight fully formed pali, with those of the lateral pairs being the largest, thickened, and roughly granulated (Figure 2C). In contrast, the pali on the dorsal and ventral directive septum are the smallest. The pali of the lateral pairs are usually taller than the septal denticles but never reach the level of the wall. Two denticles are present on each septum of the lateral pairs, mostly smaller than the palus and rarely equal (Figure 2D). The ventral and dorsal directive septum are often shorter than the lateral pairs. A distinctive feature of L1 is the occasional fusion of the ventral directive to the lateral septa of the triplet, resulting in a large palus almost equal in size to the pali of the lateral pairs (Figure 2D). Trident formation is rare, with the triplet more often free than fused (Figure 2D). Columellae are always present, sometimes as a conspicuous rod or lateral bar in the direction of the directives (Figure 2C, D). The morphological characteristics resemble those of *Porites australiensis* Vaughan, 1918, with connotations of *Porites lutea* Milne Edwards & Haime, 1851, but the absence of frequent triplet fusion or trident formation suggests a closer affinity with the former.

#### L2

The morphological characteristics of L2 differ from L1. In L2, the calices are moderately excavated, concaving downwards, with predominantly thick walls composed of three rows of denticles (Figure 2F, G). The septa are variable in thickness, ranging from thick and slightly wedge-shaped to thin and laminar (Figure 2G, H). The inner synapticular ring is mostly complete, aligning with the position of the pali. Typically, there are eight pali, with the largest and well-formed found on the lateral pairs (Figure 2G, H). However, the pali of the ventral triplet and dorsal directive are not always well developed (Figure 2H). Notably, the pali are deep-seated in the calice and either equal to or below the height of their corresponding septal denticles. Two denticles are present on each septum of the lateral pairs, typically more prominent but occasionally equal to the palus (Figure 2H). The ventral and dorsal directive septum are always shorter than the lateral pairs. The septa of the triplet are always free, with the laterals longer than the ventral septum (Figure 2G, H). Columellae are always present, mostly as a lateral bar in the direction of the directives (Figure 2G, H). The morphological characteristics of L2 closely resemble those of *Porites lobata* Dana, 1846, with a distinctive set of characters that differentiate it from L1.

#### L3

The corallite morphology observed in L3 somewhat resembles L2, with some similarities to that of L1. Within L3, the calices are moderately excavated and the walls vary, sometimes thick and not well defined or forming a ridge with rough granulated mural denticles (Figure 2J, K). Septal thickness is generally not greater than that of the interseptal loculi, and the septa are mostly characterized by a thin and laminar appearance (Figure 2L). The inner synapticular ring is more often complete, aligning with the position of the pali. Similar to L2, there are eight pali, with the largest and well-formed on the lateral pairs and the smallest and poorly developed on the ventral triplet and dorsal directive septum (Figure 2K, L). The pali are either equal to or below the height of their corresponding septal denticles, never reaching the wall. One or, more often, two denticles are present on each septum of the lateral pairs, typically more prominent but occasionally equal or slightly smaller than the palus (Figure 2L). The ventral and dorsal directive septum are always shorter than the lateral pairs. As in L2, the septa of the triplet are always free, with the laterals slightly longer or equal to the ventral septum (Figure 2K, L). Columellae are always present, as a lateral bar in the direction of the directives and occasionally as a conspicuous rod (Figure 2K, L). Structurally, L3 closely resembles *P*. *lobata*. Despite sharing some morphological similarities with *P*. *australiensis*, the less prominent pali and free margins of the triplet within L3, suggest the former.

##### Differences in polyp densities among lineages

Lineages exhibited subtle differences in mean polyp densities (Figure 6A, S7; LM: *Df* = 2, *F* = 6.586, *p* = 0.004). There was also a significant interaction between lineage and reef type (*F* = 6.903, *p* = 0.014), but reef type did not have an effect independently (*F* = 0.05, *p* = 0.819). Pairwise comparisons (based on a model without the interaction between lineage and reef type) showed that L2 colonies had ~15-20% lower mean polyp densities than L1 or L3 colonies (pairwise comparison: L1-L2 est = 0.416, *p* = 0.025; L2-L3 est = -0.570, *p* = 0.008). L1 and L3 colonies did not differ from each other (est = -0.154, *p* = 0.512). Polyp densities of L1 and L2 colonies did not significantly differ between reef types (Figure S8; L1 est = -0.416, *p* = 0.074; L2 est = -0.432, *p* = 0.073).

#### Supplementary Figures

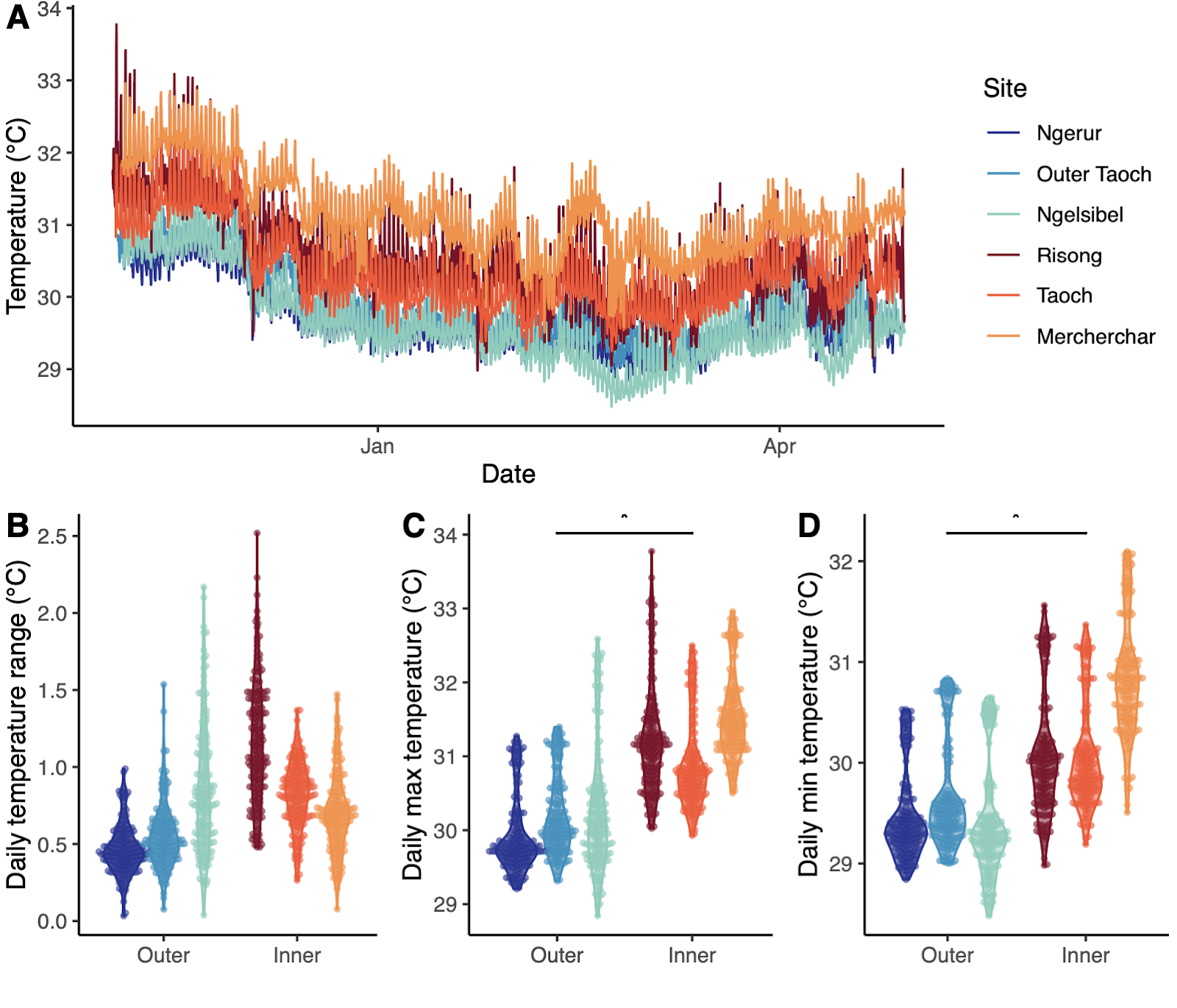

Figure S1. A) Temperature regimes differed between extreme and classic sites between November 2021 and May 2022. B) Daily temperature ranges did not differ significantly between extreme and classic sites. C) Daily maximum temperatures were 1.1 ℃ higher at extreme sites (31.2℃ ± 0.69) than classic sites (30.1℃ ± 0.66; LMM results: *F =* 25.88, *p* = 0.01). D) Daily minimum temperatures were 0.8℃ higher at extreme sites (30.3℃ ± 0.67) than classic sites (29.5℃ ± 0.52; F = 9.08, *p* = 0.04).

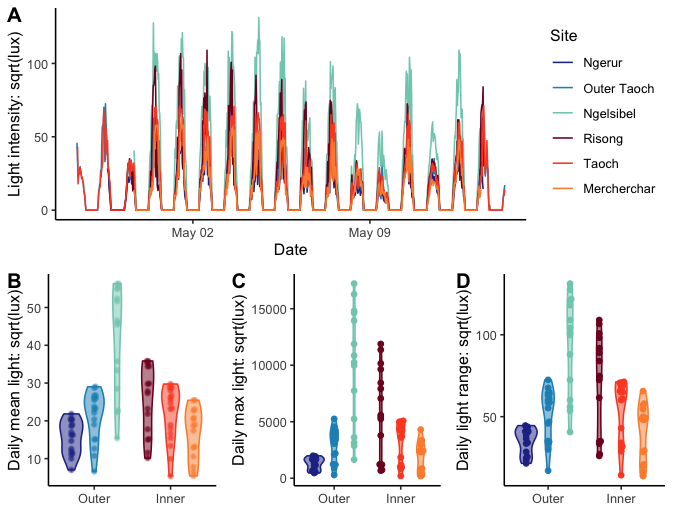

Figure S2. A) Light regimes at each of the study sites between April 28 and May 13, 2022. Mean (B; *F* = 0.337, *p* = 0.593), maximum (C; *F* = 0.068, *p* = 0.807), or daily ranges (D; *F* = 0.115, *p* = 0.752) did not differ between reef types.

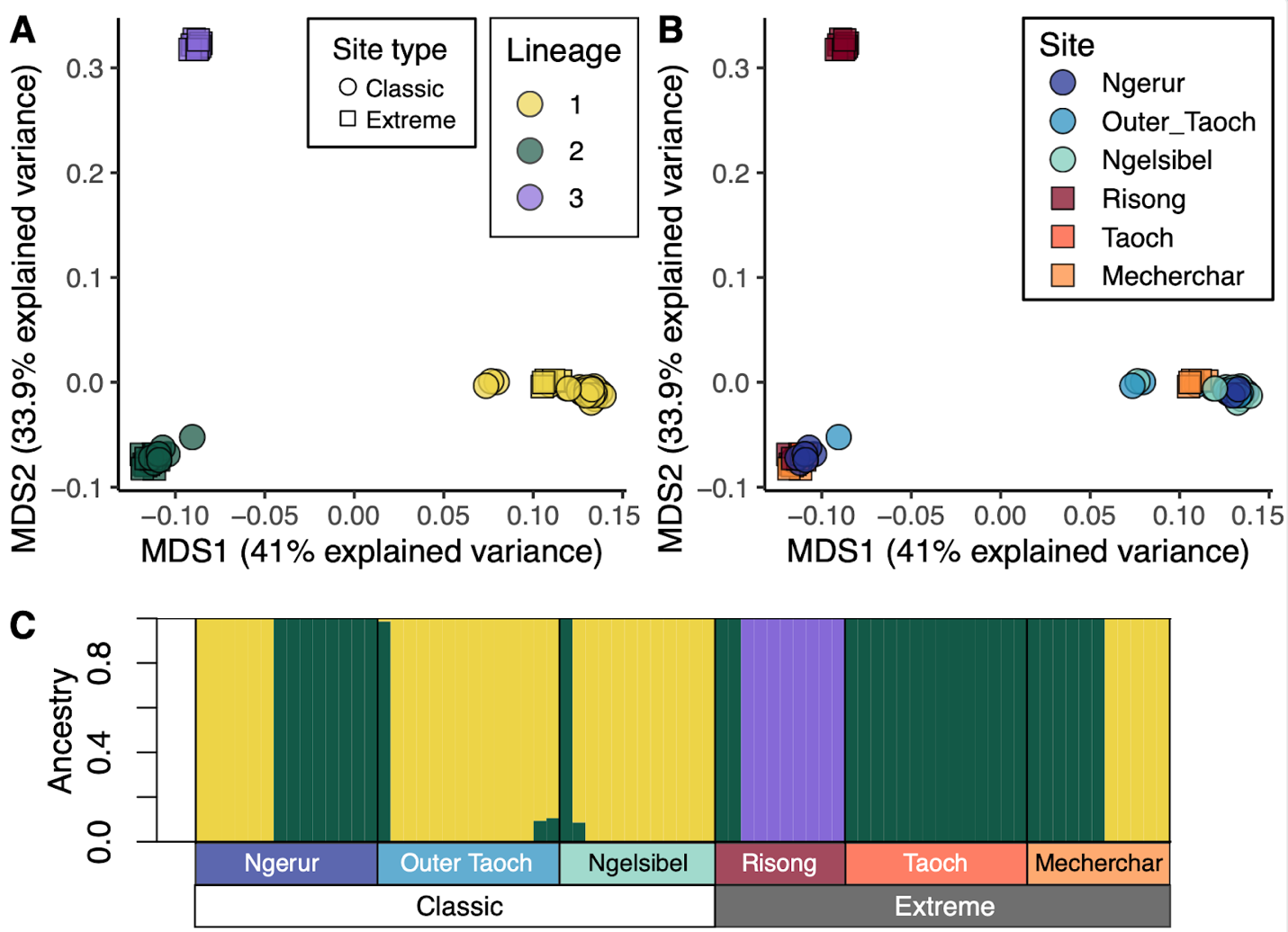

Figure S3. Population structure of massive *Porites* at six study sites in the Rock Islands of Palau based on 2b-RAD sequencing data from samples collected in November 2021 (*n* = 487,248 SNPs; see Table S1 for replication)*.* A-B) A non-metric multidimensional scaling (nMDS) plot based on genetic covariance matrices reveals three distinct genetic lineages (L1-L3). C) Results from ADMIXTURE analysis shows low, or a lack of, introgression among lineages. Each bar represents an individual coral colony. Lineage 1 (yellow) is more abundant at “classic” sites (Ngerur-Ngelsibel), whereas L2 (green) and L3 (purple) are more abundant at “extreme” sites (Risong-Mecherchar).

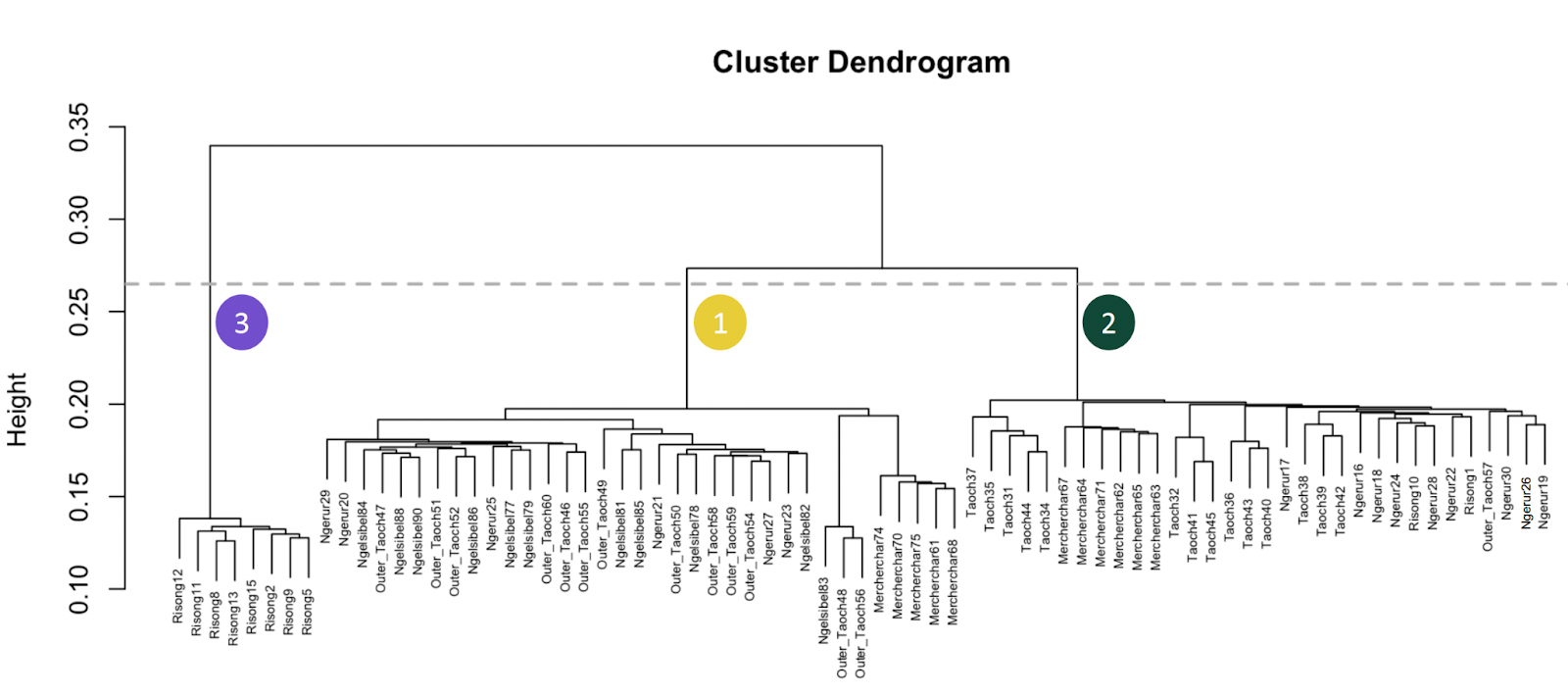

Figure S4. Cluster dendrogram of *Porites* cf. *lobata* colonies in the Rock Islands of Palau on 2b-RAD sequencing data from samples collected in November 2021 (*n* = 487,248 SNPs; see Table S1 for replication)*.* Numbers 1-3 indicate the three lineages identified in this study and colors correspond to lineage colors used throughout the manuscript.

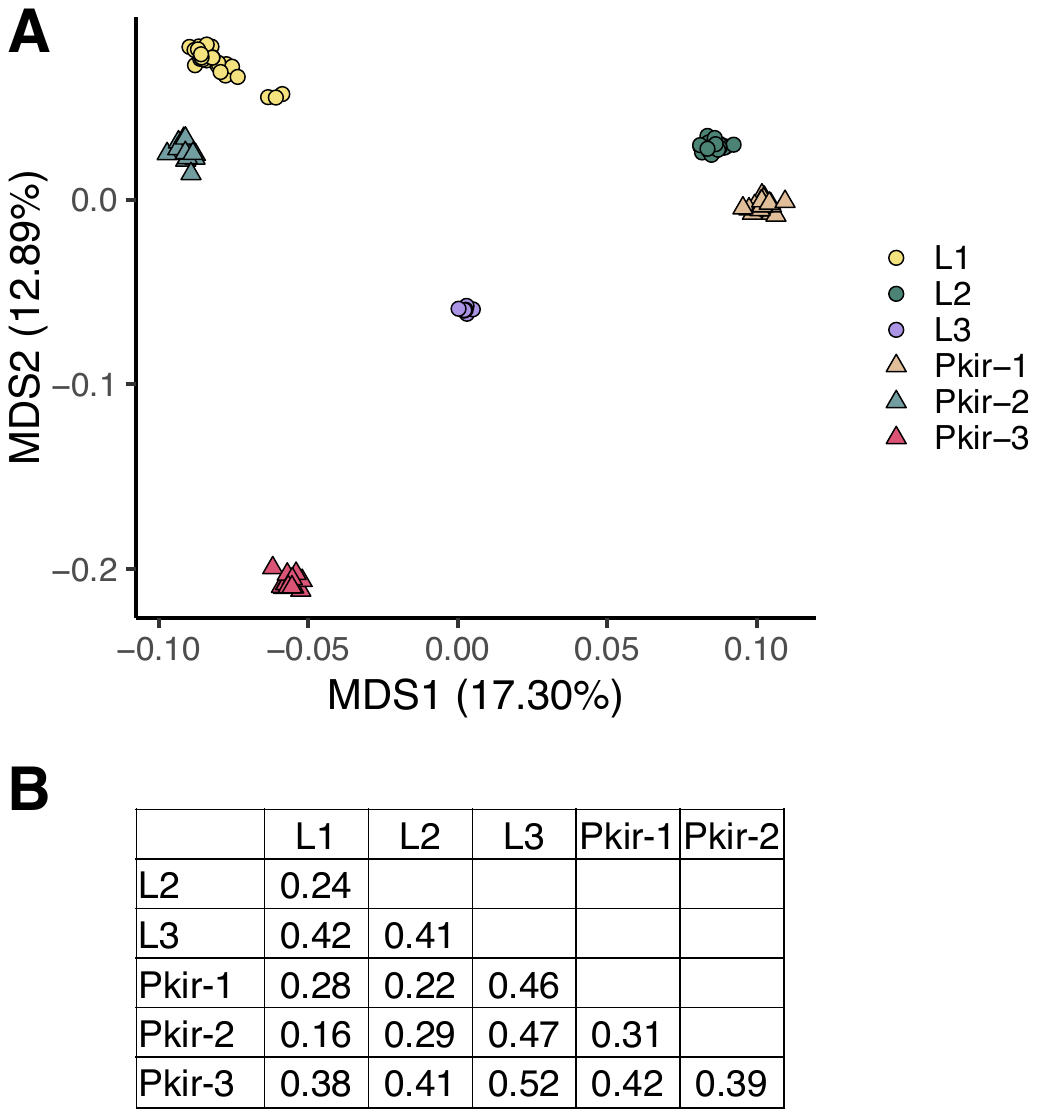

Figure S5. Population structure of cryptic lineages of massive *Porites* in Palau (L1 - L3) and Kiritimati (Pkir-1 - Pkir-3). A) non-metric multidimensional scaling (nMDS) plot showing *Porites* massive populations in Palau and Kiritimati. B) Pairwise F_st_ values show that L1 is more closely related to a lineage from Kiritimati (Pkir-2) than to other lineages in Palau. L2 is more closely related to a lineage from Kiritimati (Pkir-1) than to other lineages in Palau. L3 is highly distinct from other lineages and may be endemic to Palau. There is no lineage related to Pkir-3 in Palau. Kiritimati data adapted from Starko et al., (2023).

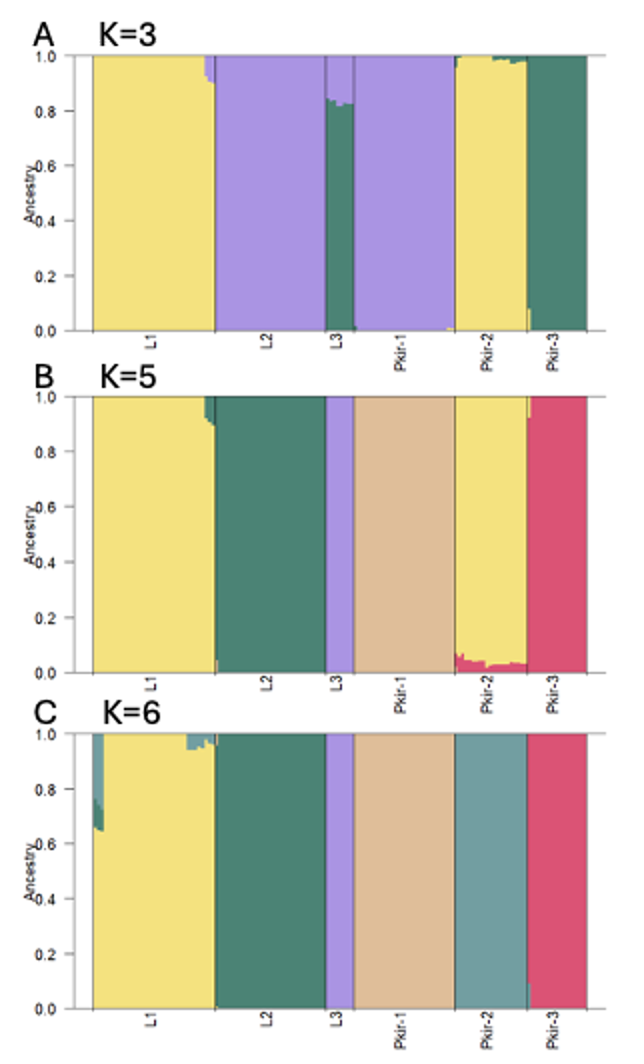

Figure S6. Results from ADMIXTURE analysis show that cryptic lineages of massive *Porites* in Palau (L1 - L3) and Kiritimati (Pkir-1 - Pkir-3) form distinct, reproductively isolated, populations. A) With K = 3, the STRUCTURE plot reveals that Palauan L1 is most closely related to Kirimatian *pkir-2*, and Palauan L2 is most closely related to Kiritimatian *pkir-1*. B) With K = 5, the plot shows that most populations are distinct, but L1 is still most closely related to pkir-2. C) With K = 6, each population forms its own cluster. Kiritimati data adapted from Starko et al., 2023.

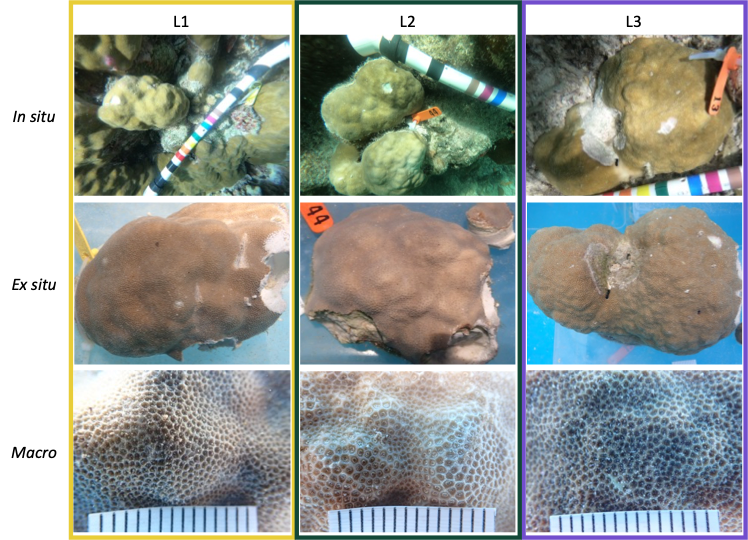

Figure S7. Representative examples of *Porites* cf. *lobata* colonies from each cryptic lineage identified in this study. Lineage 2 (L2) colonies have lower polyp densities than lineage 1 and 3 (L1, L3) colonies, and differ from L1 in terms of coloration (see Figure 2A, 2B). Scale bar in macro shots is 15 mm. Colonies displayed: 60 (L1), 44 (L2), 13 (L3).

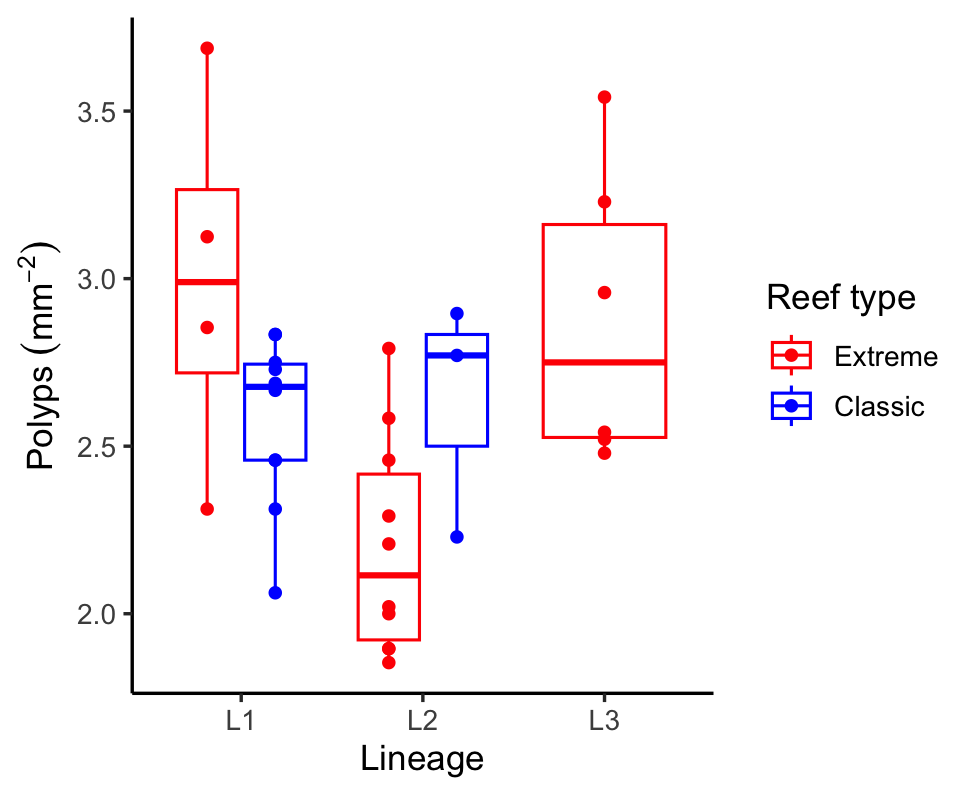

Figure S8. Polyp density in colonies from each lineage parsed by reef type.

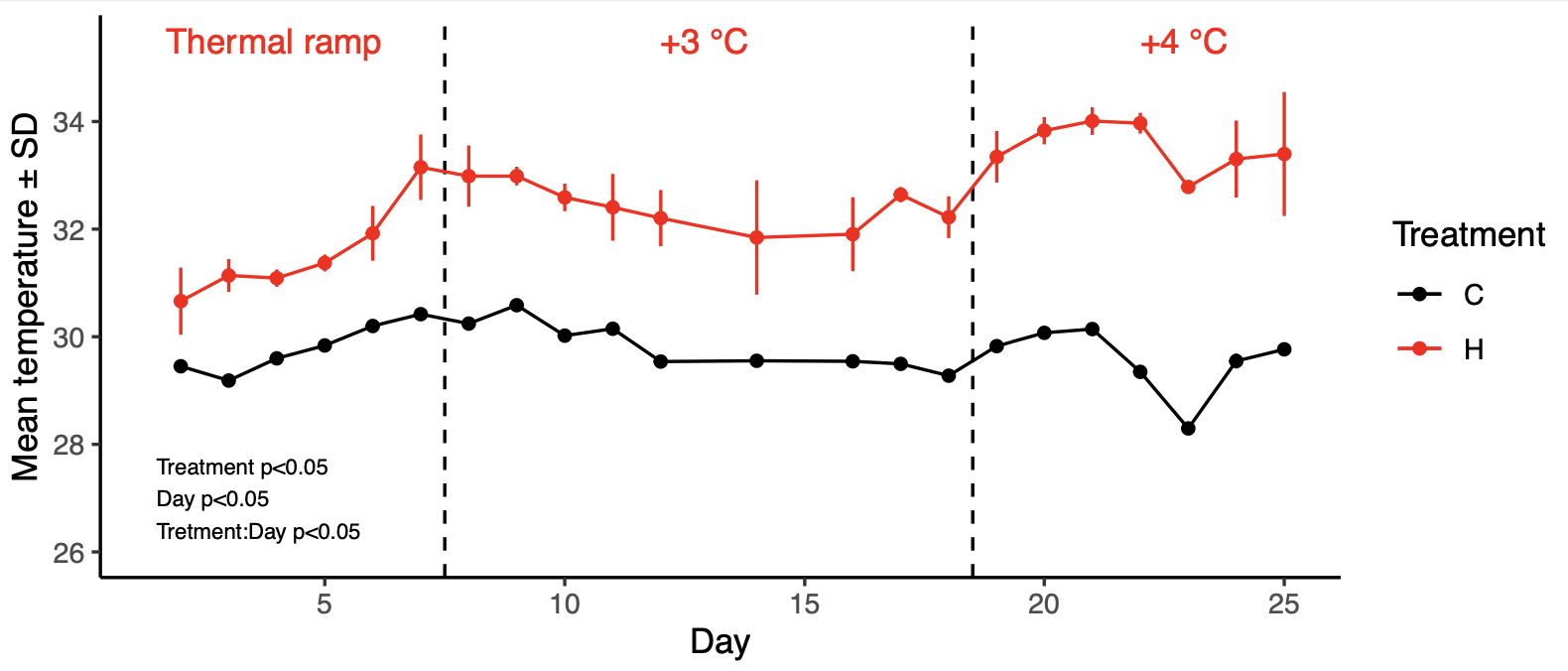

Figure S9. Daily mean temperatures in control and heat stress tanks throughout the common garden thermal challenge experiment.

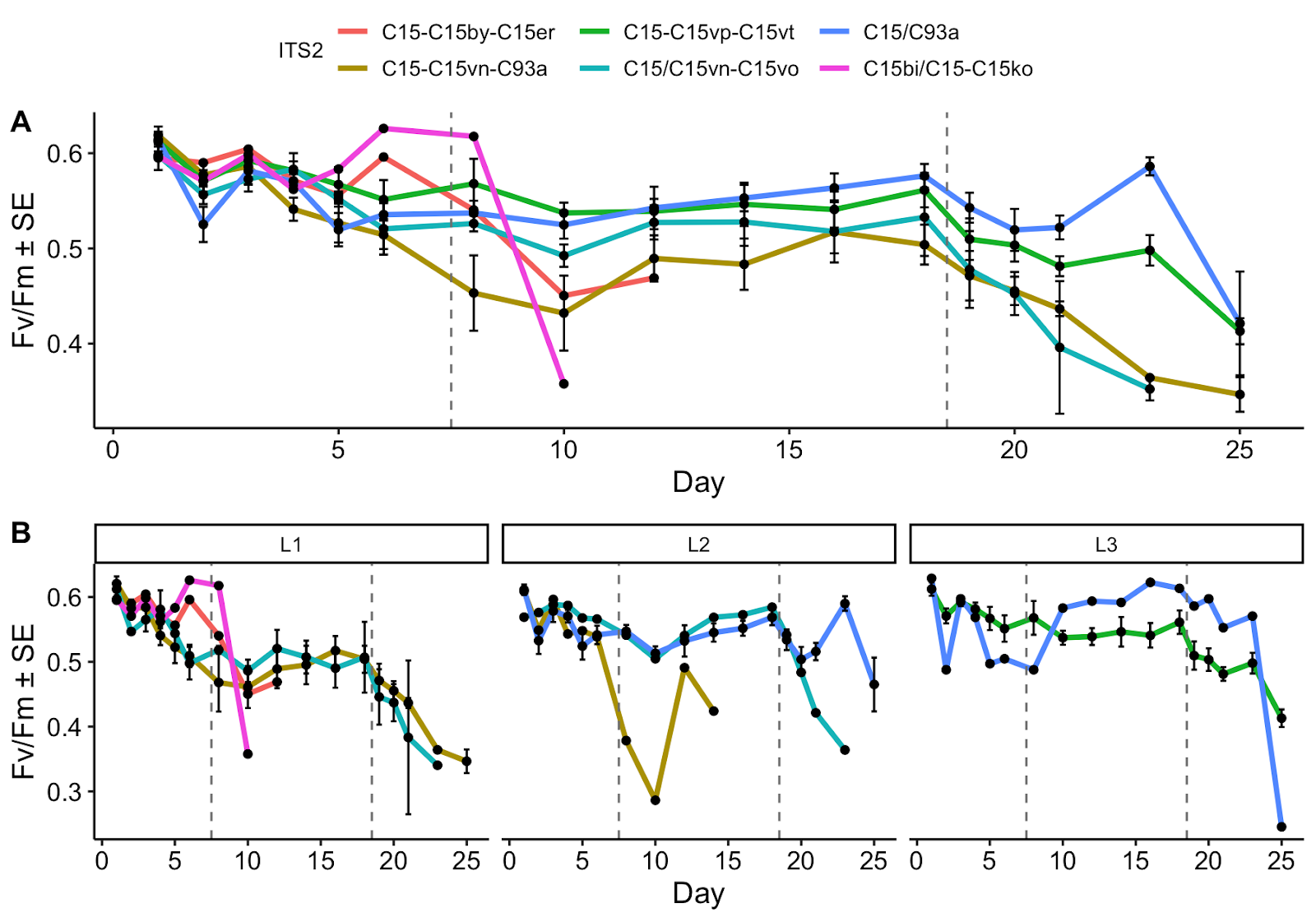

Figure S10. Photosynthetic efficiency in the thermal challenge experiment by photobiont DIV (A) and by photobiont DIV for each lineage (B).

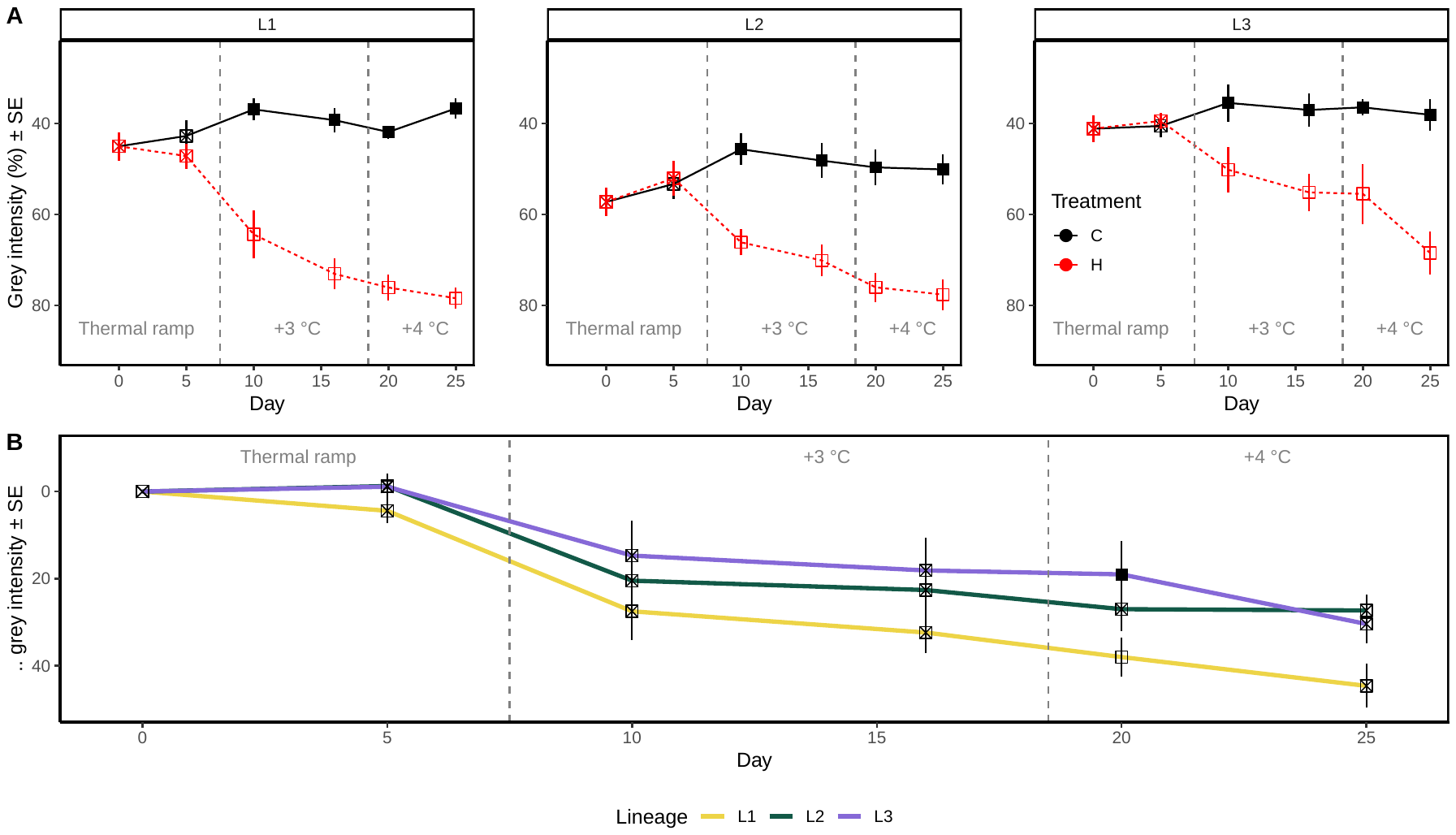

Figure S11. Paling rates differed between cryptic lineages of massive *Porites* exposed to thermal stress. A) Thermal stress caused bleaching in colonies from all three lineages. B) Paling progressed at a higher rate in L1 than L2 and L3. Closed points are significantly different from open points (*p* < 0.05; see Tables SX, SY for pairwise test results). Hatched points do not differ significantly from other points. Colony color was measured as mean intensity gray; relative coloration (panel B) was calculated by subtracting colony coloration in heat tanks from coloration in control tanks for each colony at each timepoint.

#### Supplementary Tables

Table S1. Overview of massive *Porites* samples collected for host 2b-RAD sequencing, polyp density quantification, light harvesting trait characterization, ITS2 (photobiont) and 16S (microbiome) profiling, and colonies included in the thermal challenge experiment across three classic and three extreme reefs.

| *Reef type* | *Site name* | *Total successful lineage assignments** | | | *F_st,_ PCA, cluster dendrogram, and STRUCTURE analysis* | | | *Polyp density* | | | *Light harvesting traits* | | | *Sucessful ITS2 sequencing* | | | *Successful 16S sequencing* | | | *Thermal challenge experiment* | | |
| --- | --- | --- | --- | --- | --- | --- | --- | --- | --- | --- | --- | --- | --- | --- | --- | --- | --- | --- | --- | --- | --- | --- |
|  |  | *L1* | *L2* | *L3* | *L1* | *L2* | *L3* | *L1* | *L2* | *L3* | *L1* | *L2* | *L3* | *L1* | *L2* | *L3* | *L1* | *L2* | *L3* | *L1* | *L2* | *L3* |
| *Classic* | |  |  |  |  |  |  |  |  |  |  |  |  |  |  |  |  |  |  |  |  |  |
| Ngerur | | 10 | 11 | 0 | 6 | 9 | 0 | 3 | 3 | 0 | 1 | 2 | 0 | 6 | 5 | 0 | 4 | 5 | 0 | 2 | 0 | 0 |
| Outer Taoch | | 16 | 1 | 0 | 13 | 1 | 0 | 7 | 0 | 0 | 3 | 1 | 0 | 13 | 1 | 0 | 10 | 0 | 0 | 8 | 0 | 0 |
| Ngelsibel | | 20 | 1 | 0 | 11 | 0 | 0 | 0 | 0 | 0 | 1 | 1 | 0 | 12 | 0 | 0 | 10 | 0 | 0 | 0 | 0 | 0 |
|  | **Sum** | **46** | **13** | **0** | **30** | **10** | **0** | **10** | **3** | **0** | **5** | **4** | **0** | **31** | **6** | **0** | **24** | **5** | **0** | **10** | **0** | **0** |
| *Extreme* | |  |  |  |  |  |  |  |  |  |  |  |  |  |  |  |  |  |  |  |  |  |
| Risong | | 0 | 4 | 11 | 0 | 2 | 8 | 0 | 0 | 5 | 0 | 0 | 5 | 0 | 2 | 9 | 0 | 1 | 5 | 0 | 1 | 4 |
| Taoch | | 0 | 16 | 0 | 0 | 14 | 0 | 0 | 5 | 0 | 0 | 5 | 0 | 0 | 14 | 0 | 0 | 5 | 0 | 0 | 6 | 0 |
| Mecherchar | | 5 | 6 | 3 | 5 | 6 | 0 | 4 | 5 | 1 | 0 | 1 | 0 | 4 | 6 | 1 | 3 | 5 | 0 | 0 | 2 | 1 |
|  | **Sum** | **5** | **26** | **14** | **5** | **22** | **8** | **4** | **10** | **6** | **0** | **6** | **5** | **4** | **22** | **10** | **3** | **11** | **5** | **0** | **9** | **5** |
|  | **Total** | **51** | **39** | **14** | **35** | **32** | **8** | **14** | **13** | **6** | **5** | **10** | **5** | **35** | **28** | **10** | **27** | **16** | **5** | **10** | **9** | **5** |

*Total numbers combined after three 2b-RAD runs

Table S2. Pairwise comparisons of Fv/Fm values between control and heat stress treatments for each tested lineage. Pairwise tests were conducted using the package emmeans with a Tukey correction and constrained to tests between control and heat treatment colonies for each lineage for each individual day.

| *Day* | *Lineage* | | | | | |
| --- | --- | --- | --- | --- | --- | --- |
|  | *L1* | | *L2* | | *L3* | |
|  | *Estimate* | *p* | *Estimate* | *p* | *Estimate* | *p* |
| 1 | -0.005 | 0.771 | 0.010 | 0.593 | -0.008 | 0.732 |
| 2 | 0.015 | 0.388 | 0.044 | **0.025** | 0.039 | 0.093 |
| 3 | 0.027 | 0.123 | 0.038 | 0.051 | 0.022 | 0.351 |
| 4 | 0.045 | **0.010** | 0.039 | **0.047** | 0.036 | 0.125 |
| 5 | 0.045 | **0.009** | 0.054 | **0.006** | 0.042 | 0.073 |
| 6 | 0.063 | **0.000** | 0.052 | **0.008** | 0.063 | **0.007** |
| 8 | 0.095 | **<.0001** | 0.088 | **<.0001** | 0.056 | **0.017** |
| 10 | 0.144 | **<.0001** | 0.107 | **<.0001** | 0.060 | **0.010** |
| 12 | 0.113 | **<.0001** | 0.070 | **0.000** | 0.060 | **0.010** |
| 14 | 0.120 | **<.0001** | 0.081 | **<.0001** | 0.091 | **0.000** |
| 16 | 0.106 | **<.0001** | 0.056 | **0.006** | 0.074 | **0.002** |
| 18 | 0.113 | **<.0001** | 0.041 | **0.047** | 0.055 | **0.019** |
| 19 | 0.149 | **<.0001** | 0.064 | **0.002** | 0.085 | **0.000** |
| 20 | 0.158 | **<.0001** | 0.106 | **<.0001** | 0.087 | **0.000** |
| 21 | 0.175 | **<.0001** | 0.105 | **<.0001** | 0.128 | **<.0001** |
| 23 | 0.244 | **<.0001** | 0.067 | **0.002** | 0.125 | **<.0001** |
| 25 | 0.252 | **<.0001** | 0.138 | **<.0001** | 0.249 | **<.0001** |

Table S3. Pairwise comparisons of Fv/Fm values between lineages within the heat treatment. Pairwise comparisons were conducted using the package emmeans with a false discovery rate correction and constrained to tests between lineages for each individual day.

| Day | Treatment | Comparison | Estimate | SE | Df | t.ratio | *p* |
| --- | --- | --- | --- | --- | --- | --- | --- |
| 1 | C | L1 - L2 | -1.1E-02 | 0.02 | 558 | -0.65 | 0.929 |
| 1 | C | L1 - L3 | -3.3E-03 | 0.02 | 558 | -0.16 | 0.945 |
| 1 | C | L2 - L3 | 7.9E-03 | 0.02 | 558 | 0.38 | 0.945 |
| 2 | C | L1 - L2 | -2.0E-03 | 0.02 | 558 | -0.12 | 0.945 |
| 2 | C | L1 - L3 | -5.2E-03 | 0.02 | 558 | -0.25 | 0.945 |
| 2 | C | L2 - L3 | -3.2E-03 | 0.02 | 558 | -0.15 | 0.945 |
| 3 | C | L1 - L2 | -1.1E-02 | 0.02 | 558 | -0.63 | 0.929 |
| 3 | C | L1 - L3 | -2.7E-03 | 0.02 | 558 | -0.13 | 0.945 |
| 3 | C | L2 - L3 | 8.1E-03 | 0.02 | 558 | 0.39 | 0.945 |
| 4 | C | L1 - L2 | -6.3E-03 | 0.02 | 558 | -0.36 | 0.945 |
| 4 | C | L1 - L3 | -1.3E-02 | 0.02 | 558 | -0.62 | 0.929 |
| 4 | C | L2 - L3 | -6.5E-03 | 0.02 | 558 | -0.31 | 0.945 |
| 5 | C | L1 - L2 | -9.3E-03 | 0.02 | 558 | -0.54 | 0.930 |
| 5 | C | L1 - L3 | -1.2E-02 | 0.02 | 558 | -0.59 | 0.929 |
| 5 | C | L2 - L3 | -2.8E-03 | 0.02 | 558 | -0.13 | 0.945 |
| 6 | C | L1 - L2 | -5.9E-04 | 0.02 | 558 | -0.03 | 0.982 |
| 6 | C | L1 - L3 | -1.2E-02 | 0.02 | 558 | -0.58 | 0.929 |
| 6 | C | L2 - L3 | -1.1E-02 | 0.02 | 558 | -0.54 | 0.930 |
| 8 | C | L1 - L2 | -1.0E-02 | 0.02 | 558 | -0.58 | 0.929 |
| 8 | C | L1 - L3 | -5.8E-03 | 0.02 | 558 | -0.28 | 0.945 |
| 8 | C | L2 - L3 | 4.2E-03 | 0.02 | 558 | 0.20 | 0.945 |
| 10 | C | L1 - L2 | 6.6E-03 | 0.02 | 558 | 0.38 | 0.945 |
| 10 | C | L1 - L3 | -8.5E-03 | 0.02 | 558 | -0.41 | 0.945 |
| 10 | C | L2 - L3 | -1.5E-02 | 0.02 | 558 | -0.72 | 0.929 |
| 12 | C | L1 - L2 | 2.2E-03 | 0.02 | 558 | 0.13 | 0.945 |
| 12 | C | L1 - L3 | -7.4E-03 | 0.02 | 558 | -0.36 | 0.945 |
| 12 | C | L2 - L3 | -9.6E-03 | 0.02 | 558 | -0.46 | 0.945 |
| 14 | C | L1 - L2 | 4.7E-03 | 0.02 | 558 | 0.27 | 0.945 |
| 14 | C | L1 - L3 | -3.1E-02 | 0.02 | 558 | -1.53 | 0.520 |
| 14 | C | L2 - L3 | -3.6E-02 | 0.02 | 558 | -1.72 | 0.414 |
| 16 | C | L1 - L2 | 2.4E-03 | 0.02 | 558 | 0.14 | 0.945 |
| 16 | C | L1 - L3 | -1.8E-02 | 0.02 | 558 | -0.88 | 0.901 |
| 16 | C | L2 - L3 | -2.1E-02 | 0.02 | 558 | -0.98 | 0.842 |
| 18 | C | L1 - L2 | 4.1E-03 | 0.02 | 558 | 0.24 | 0.945 |
| 18 | C | L1 - L3 | -7.6E-03 | 0.02 | 558 | -0.37 | 0.945 |
| 18 | C | L2 - L3 | -1.2E-02 | 0.02 | 558 | -0.56 | 0.930 |
| 19 | C | L1 - L2 | 1.4E-02 | 0.02 | 558 | 0.83 | 0.911 |
| 19 | C | L1 - L3 | 1.7E-03 | 0.02 | 558 | 0.08 | 0.963 |
| 19 | C | L2 - L3 | -1.3E-02 | 0.02 | 558 | -0.60 | 0.929 |
| 20 | C | L1 - L2 | -8.1E-05 | 0.02 | 558 | -0.01 | 0.996 |
| 20 | C | L1 - L3 | -4.9E-03 | 0.02 | 558 | -0.24 | 0.945 |
| 20 | C | L2 - L3 | -4.9E-03 | 0.02 | 558 | -0.23 | 0.945 |
| 21 | C | L1 - L2 | -3.9E-03 | 0.02 | 558 | -0.23 | 0.945 |
| 21 | C | L1 - L3 | -2.4E-02 | 0.02 | 558 | -1.17 | 0.755 |
| 21 | C | L2 - L3 | -2.0E-02 | 0.02 | 558 | -0.96 | 0.842 |
| 23 | C | L1 - L2 | -9.0E-03 | 0.02 | 558 | -0.52 | 0.932 |
| 23 | C | L1 - L3 | -3.3E-02 | 0.02 | 558 | -1.59 | 0.512 |
| 23 | C | L2 - L3 | -2.4E-02 | 0.02 | 558 | -1.14 | 0.755 |
| 25 | C | L1 - L2 | 1.1E-02 | 0.02 | 558 | 0.66 | 0.929 |
| 25 | C | L1 - L3 | -2.0E-02 | 0.02 | 558 | -0.98 | 0.842 |
| 25 | C | L2 - L3 | -3.2E-02 | 0.02 | 558 | -1.50 | 0.524 |
| 1 | H | L1 - L2 | 1.3E-02 | 0.02 | 558 | 0.77 | 0.929 |
| 1 | H | L1 - L3 | -1.3E-03 | 0.02 | 558 | -0.07 | 0.967 |
| 1 | H | L2 - L3 | -1.5E-02 | 0.02 | 558 | -0.69 | 0.929 |
| 2 | H | L1 - L2 | 2.2E-02 | 0.02 | 558 | 1.30 | 0.688 |
| 2 | H | L1 - L3 | 1.9E-02 | 0.02 | 558 | 0.91 | 0.882 |
| 2 | H | L2 - L3 | -3.6E-03 | 0.02 | 558 | -0.17 | 0.945 |
| 3 | H | L1 - L2 | -4.2E-03 | 0.02 | 558 | -0.24 | 0.945 |
| 3 | H | L1 - L3 | -1.2E-02 | 0.02 | 558 | -0.58 | 0.929 |
| 3 | H | L2 - L3 | -7.9E-03 | 0.02 | 558 | -0.38 | 0.945 |
| 4 | H | L1 - L2 | -1.5E-02 | 0.02 | 558 | -0.86 | 0.904 |
| 4 | H | L1 - L3 | -2.4E-02 | 0.02 | 558 | -1.14 | 0.755 |
| 4 | H | L2 - L3 | -8.7E-03 | 0.02 | 558 | -0.41 | 0.945 |
| 5 | H | L1 - L2 | 1.3E-02 | 0.02 | 558 | 0.75 | 0.929 |
| 5 | H | L1 - L3 | -1.6E-02 | 0.02 | 558 | -0.80 | 0.925 |
| 5 | H | L2 - L3 | -2.9E-02 | 0.02 | 558 | -1.40 | 0.615 |
| 6 | H | L1 - L2 | -1.9E-02 | 0.02 | 558 | -1.07 | 0.802 |
| 6 | H | L1 - L3 | -1.4E-02 | 0.02 | 558 | -0.70 | 0.929 |
| 6 | H | L2 - L3 | 4.2E-03 | 0.02 | 558 | 0.20 | 0.945 |
| 8 | H | L1 - L2 | -2.2E-02 | 0.02 | 558 | -1.25 | 0.695 |
| 8 | H | L1 - L3 | -4.5E-02 | 0.02 | 558 | -2.20 | 0.169 |
| 8 | H | L2 - L3 | -2.4E-02 | 0.02 | 558 | -1.13 | 0.755 |
| 10 | H | L1 - L2 | -2.7E-02 | 0.02 | 558 | -1.58 | 0.512 |
| 10 | H | L1 - L3 | -9.1E-02 | 0.02 | 558 | -4.42 | **0.000** |
| 10 | H | L2 - L3 | -6.4E-02 | 0.02 | 558 | -3.04 | **0.033** |
| 12 | H | L1 - L2 | -3.7E-02 | 0.02 | 569 | -2.08 | 0.217 |
| 12 | H | L1 - L3 | -5.7E-02 | 0.02 | 566 | -2.71 | **0.047** |
| 12 | H | L2 - L3 | -2.0E-02 | 0.02 | 558 | -0.96 | 0.842 |
| 14 | H | L1 - L2 | -3.6E-02 | 0.02 | 581 | -1.97 | 0.263 |
| 14 | H | L1 - L3 | -6.3E-02 | 0.02 | 575 | -2.97 | **0.036** |
| 14 | H | L2 - L3 | -2.8E-02 | 0.02 | 558 | -1.31 | 0.688 |
| 16 | H | L1 - L2 | -2.4E-02 | 0.02 | 602 | -1.26 | 0.695 |
| 16 | H | L1 - L3 | -4.2E-02 | 0.02 | 585 | -1.91 | 0.288 |
| 16 | H | L2 - L3 | -1.8E-02 | 0.02 | 568 | -0.82 | 0.911 |
| 18 | H | L1 - L2 | -5.9E-02 | 0.02 | 602 | -3.02 | **0.033** |
| 18 | H | L1 - L3 | -6.1E-02 | 0.02 | 585 | -2.80 | **0.045** |
| 18 | H | L2 - L3 | -3.0E-03 | 0.02 | 568 | -0.14 | 0.945 |
| 19 | H | L1 - L2 | -5.2E-02 | 0.02 | 602 | -2.69 | **0.047** |
| 19 | H | L1 - L3 | -6.3E-02 | 0.02 | 585 | -2.86 | **0.042** |
| 19 | H | L2 - L3 | -1.1E-02 | 0.02 | 568 | -0.50 | 0.945 |
| 20 | H | L1 - L2 | -5.2E-02 | 0.02 | 602 | -2.70 | **0.047** |
| 20 | H | L1 - L3 | -8.5E-02 | 0.02 | 585 | -3.87 | **0.003** |
| 20 | H | L2 - L3 | -3.3E-02 | 0.02 | 568 | -1.53 | 0.520 |
| 21 | H | L1 - L2 | -7.3E-02 | 0.02 | 622 | -3.53 | **0.009** |
| 21 | H | L1 - L3 | -7.6E-02 | 0.02 | 597 | -3.35 | **0.015** |
| 21 | H | L2 - L3 | -2.7E-03 | 0.02 | 578 | -0.12 | 0.945 |
| 23 | H | L1 - L2 | -1.7E-01 | 0.03 | 658 | -6.62 | **<.0001** |
| 23 | H | L1 - L3 | -1.5E-01 | 0.03 | 641 | -5.51 | **<.0001** |
| 23 | H | L2 - L3 | 2.4E-02 | 0.02 | 591 | 1.04 | 0.828 |
| 25 | H | L1 - L2 | -8.8E-02 | 0.03 | 665 | -2.85 | **0.042** |
| 25 | H | L1 - L3 | -2.3E-02 | 0.03 | 657 | -0.74 | 0.929 |
| 25 | H | L2 - L3 | 6.5E-02 | 0.02 | 605 | 2.75 | **0.047** |

Table S4. Pairwise comparisons of color (mean intensity gray %) values between control and heat stress treatments for each tested lineage. Pairwise tests were conducted using the package emmeans with a false discovery rate correction and constrained to tests between control and heat treatment colonies for each lineage for each individual day.

| *Day* | *Lineage* | | | | | |
| --- | --- | --- | --- | --- | --- | --- |
|  | *L1* | | *L2* | | *L3* | |
|  | *Estimate* | *p* | *Estimate* | *p* | *Estimate* | *p* |
| 0 | 0.000 | 1.000 | 0.000 | 1.000 | 0.000 | 1.000 |
| 5 | -4.42 | 0.367 | 1.26 | 0.981 | 1.11 | 1.000 |
| 10 | -27.50 | **<.0001** | -20.47 | **<0.0001** | -14.71 | **0.014** |
| 16 | -34.66 | **<.0001** | -22.34 | **<0.0001** | -18.12 | **0.002** |
| 20 | -35.83 | **<.0001** | -26.77 | **<0.0001** | -19.02 | **0.001** |
| 25 | -43.89 | **<.0001** | -27.65 | **<0.0001** | -30.41 | **<.0001** |

Table S5. Pairwise comparisons of relative color values (mean intensity gray in heat - control colonies) between lineages. Pairwise comparisons were conducted using the package emmeans with a false discovery rate correction and constrained to tests between lineages for each individual day.

| *Day* | *Lineage* | | | | | |
| --- | --- | --- | --- | --- | --- | --- |
|  | *L1-L2* | | *L1-L3* | | *L2-L3* | |
|  | *Estimate* | *p* | *Estimate* | *p* | *Estimate* | *p* |
| 0 | 0.00 | 1.00 | 0.00 | 1.00 | 0.00 | 1.00 |
| 5 | 5.68 | 0.88 | 5.53 | 1.00 | -0.15 | 1.00 |
| 10 | 7.04 | 0.58 | 12.80 | 0.14 | 5.76 | 1.00 |
| 16 | 9.95 | 0.29 | 14.54 | 0.10 | 4.59 | 1.00 |
| 20 | 11.26 | 0.33 | 19.34 | **0.04** | 8.08 | 0.68 |
| 25 | 17.56 | 0.08 | 15.57 | 0.19 | -1.99 | 1.00 |
